# Myosin VI regulates ciliogenesis by promoting the turnover of the centrosomal/satellite protein OFD1

**DOI:** 10.1101/2021.06.18.448975

**Authors:** Elisa Magistrati, Giorgia Maestrini, Mariana Lince-Faria, Galina Beznoussenko, Alexandre Mironov, Elena Maspero, Mónica Bettencourt-Dias, Simona Polo

**Affiliations:** IFOM, Fondazione Istituto FIRC di Oncologia Molecolare, Milan, Italy; IGC, Instituto Gulbenkian de Ciencia, Oeiras, Portugal; Dipartimento di Oncologia ed Emato-oncologia, Università degli Studi di Milano, Milan, Italy

## Abstract

The actin motor protein myosin VI is a multivalent protein with diverse functions. Here, we identified and characterised a myosin VI ubiquitous interactor, the oral-facial-digital syndrome 1 (OFD1) protein, whose mutations cause malformations of the face, oral cavity, digits, and polycystic kidney disease. We found that myosin VI regulates the localisation of OFD1 at the centrioles and, as a consequence, the recruitment of the distal appendage protein cep164. Myosin VI depletion in non-tumoural cell lines causes an aberrant localisation of OFD1 along the centriolar walls, which is due to a reduction in the OFD1 mobile fraction. Finally, loss of myosin VI triggers a severe defect in ciliogenesis that could be causally linked to an impairment in the autophagic removal of OFD1 from satellites. Altogether, our results highlight an unprecedent layer of regulation of OFD1 and a pivotal role of myosin VI in coordinating the formation of the distal appendages and primary cilium with important implications for the genetic disorders known as ciliopathies.

## Introduction

Primary cilia are sensory structures extending from the surface of mammalian cells, with a major role in several signalling pathways essential for growth and differentiation, such as the Hedgehog and the Wnt signalling pathways (Berbari et al. 2009; Malicki and Johnson 2017). The formation of the primary cilium is a highly regulated multi-step process that occurs in cells that exit the cell cycle and become quiescent. The primary cilium originates from the older centriole, called mother centriole, which docks to the plasma membrane and acts as basal body for the assembly of the microtubule ciliary axoneme (Sanchez and Dynlacht 2016). Impairment in the formation or function of the primary cilia leads to a variety of severe genetic syndromes, termed ciliopathies (Reiter and Leroux 2017). This group of disorders includes the oral-facial-digital (OFD) type I, the Simpson–Golabi–Behmel type 2 and the Joubert syndromes, which are caused by mutations in the OFD1 gene (Ferrante et al. 2006; Coene et al. 2009; Ferrante et al. 2001; Zullo et al. 2010; Field et al. 2012; Feather et al. 1997; Budny et al. 2006). The mechanisms underlying many of the disease phenotypes associated with ciliary dysfunction have yet to be fully elucidated. Dissecting the regulatory mechanisms of OFD1 has the potential to offer new therapeutic tools for the treatment of ciliopathies.

OFD1 is a protein component of the centrioles and pericentriolar satellites, and acts both as a positive and negative regulator of primary ciliogenesis (Ferrante et al. 2006; Tang et al. 2013; Morleo and Franco 2020). The localisation of OFD1 is differentially regulated depending on the cell compartment. At the centrioles, OFD1 is recruited to the distal tip through the interaction with the C2 domain containing 3 centriole elongation regulator (C2CD3), a protein required for centriole elongation (Thauvin-Robinet et al. 2014). OFD1 recruitment is under the control of a subset of centriole proteins that are present only in the daughter centriole (daughter centriole-specific proteins), but the molecular link is unknown (Wang et al. 2018). OFD1 in turn promotes the recruitment of the distal appendages, structures present on the distal side of the mother centriole. These structures are essential for the docking of the mother centriole to the cellular membrane and its conversion to basal body, required for ciliogenesis (Singla et al. 2010; Wang et al. 2018). Furthermore, OFD1 is required for the recruitment of IFT88, a protein essential for the assembly of the primary cilium (Singla et al. 2010). Despite the growing literature that explores the process of primary cilium formation, the mechanistic details of this sequential recruitment of C2CD3, OFD1 and distal appendages remain unclear.

OFD1 is also localised at the centriolar satellites, which are non-membrane electron-dense particles containing regulatory proteins of centrosomes and cilia (Lopes et al. 2011). This localisation depends on the trafficking protein particle complex subunit 8 (TRAPPC8), which mediates the interaction between OFD1 and the main structural component of the centriolar satellites, namely PCM1 (Zhang et al. 2020).

While the centriolar pool of OFD1 is rather stable, the satellite pool shows a higher turnover (Tang et al. 2013). Indeed, upon serum starvation, OFD1 is removed from the satellites by autophagy, and this process is required for ciliogenesis (Tang et al. 2013; Park et al. 2018). Interestingly, the depletion of OFD1 from the satellites is sufficient to induce ciliogenesis also in autophagy-impaired cells (Tang et al. 2013), indicating that OFD1 turnover at the satellites controls *per se* the formation of the primary cilium. Moreover, OFD1 appears to regulate autophagosome biogenesis in a feedback loop that aims at limiting autophagy activation (Morleo et al. 2020).

A proteomic study identified OFD1 as a possible interactor of myosin VI (O’Loughlin, Masters, and Buss 2018), a unique motor protein that moves towards the minus end of the actin filaments (Magistrati and Polo 2020). The molecular pathways and the physiological relevance of this interaction remain unknown. Here, we characterized the interaction between the two proteins and we investigated the role of myosin VI in OFD1 regulation.

## Results

### OFD1 interacts with both myosin VI isoforms

The functional and phenotypic diversity associated with myosin VI arises from alternatively-spliced isoforms that interact with multiple proteins (Magistrati and Polo 2020). To capture common interactors, we used co-immunoprecipation assay and analysed by mass spectrometry the interactome of endogenous myosin VI in a set of cell lines of epithelial origin expressing the different isoforms at various level, namely MDA-MB-231, HeLa, MCF10A, and Caco-2 cells (Supplementary Table 1). 13 proteins were found in all four cell lines (Figure 1A) and most of them were identified as myosin VI interactors for the first time. We focused our attention on OFD1 and we validated the interaction between myosin VI and OFD1, using a second anti-myosin VI antibody (Figure 1B) and by performing a reverse co-immunoprecipitation experiment (Figure 1C).

**Figure 1:**
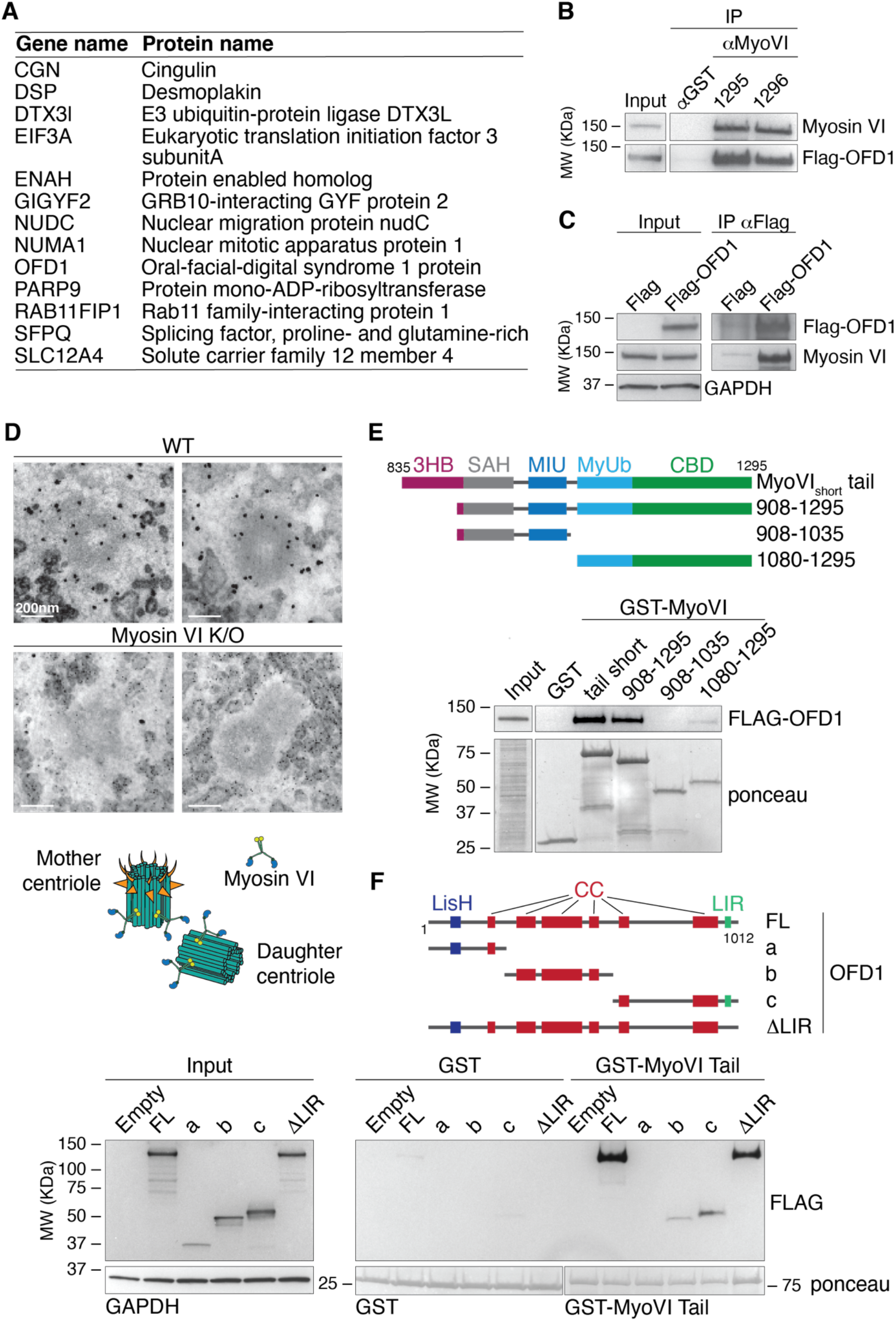
OFD1 interacts with both myosin VI isoforms. (A) List of myosin VI interactors found in the four cell lines tested. See Supplementary Table 1 for the full list of interactors. (B) Total lysates from HEK293T transfected with Flag-OFD1 were IP with anti-myosin VI antibodies (1295 and 1296) and an unrelated rabbit antibody (anti-GST, as control). IB was performed with anti-Flag and anti-myosin VI antibodies. (C) Total lysates from HEK293T transfected with Flag-OFD1 or Flag (as control) were IP with anti-Flag antibody-conjugated beads. IB was performed with anti-Flag and anti-myosin VI antibodies. (D) Selected sections deriving from TEM analysis of A549 wild-type versus myosin VI KO cells. The cells were immunogold-labelled with anti-myosin VI antibody. Scale bar, 200 nm. Bottom, representation of the estimated localisation of myosin VI at the centrioles. (E) GST pulldown assay using the indicated myosin VI constructs or GST alone (as control) and lysates from HEK293T cells transfected with Flag-OFD1 construct. IB was performed with anti-Flag antibody. Ponceau staining as indicated. (F) GST pulldown assay using myosin VI tail (835-1295) construct (or GST alone as control) and lysates from HEK293T cells transfected with the indicated Flag-OFD1 constructs (or Flag alone, as control). IB was performed with anti-Flag antibody. Ponceau staining as indicated.

Since OFD1 is a centriolar protein, we assessed the localisation of myosin VI in this organelle. By immunofluorescence (Supplementary Figure 1A) and proximity ligation assay (PLA, Supplementary Figure 1B) we could demonstrate that myosin VI localises at centrioles in correspondence of OFD1 staining, suggesting that the interaction between the two proteins can occur at least at the level of the centrioles. To further confirm this idea, we performed immunogold electron microscopy analysis of the endogenous myosin VI. As expected, myosin VI shows multiple signals corresponding to the various cytoplasmatic organelles where it performs its function (Magistrati and Polo 2020). Notably, a specific signal was evident at the level of centriole walls in wild-type but not in a myosin VI knock-out (KO) cell line (Figure 1D).

Next, we examined in detail the interaction between myosin VI and OFD1. Through immunoprecipitation and pull-down experiments, we confirmed that the short and long myosin VI isoforms can equally bind OFD1 and that the binding is mediated by the tail domain of myosin VI (Supplementary Figure 2A, B). In an attempt to characterise the site of the interaction, we focused on the major, well-characterised myosin VI cargo binding sites, namely the WWY motif (Spudich et al. 2007), the RRL motif (Sahlender et al. 2005) and the MyUb (Myosin VI Ubiquitin-binding) domain (Penengo et al. 2006; He et al. 2016). Point mutations in key residues did not affect the binding to OFD1 (Supplementary Figure 2C, D), prompting us to look for another binding region inside the myosin VI tail. By structure-function analysis we identified the C-terminal region that starts from the MyUb as critical, but not sufficient to reach the OFD1 binding level of the entire tail domain (Figure 1E). Interestingly, the SAH domain, while it does not interact with OFD1 *per se* (Supplementary Figure 2E), appears to be required for the maximum binding (Figure 1E), implying that the conformation of the tail may be important for the interaction (Magistrati and Polo 2020).

We then investigated the region of OFD1 that mediates myosin VI binding. Starting from the N-terminus, OFD1 is composed of a highly conserved Lis1 homology (LisH) motif that is important for protein-protein interactions, protein stability and intracellular localisation (Singla et al. 2010), followed by six coiled-coils that are important for centrosomal localisation (Romio et al. 2004; Lopes et al. 2011) and a C-terminal LIR domain that mediates the binding to the autophagosomes (Morleo et al. 2020). Pull-down experiments showed that OFD1 binds myosin VI through its coiled-coils region, while the LIR domain and the N-terminal region containing the LisH motif appear dispensable for the interaction (Figure 1F). Unfortunately, the bad behaviour of the shorter constructs of OFD1 precluded further analysis of the interaction boundaries, as well as possible crystallisation attempts.

### Myosin VI regulates OFD1 levels at the centrioles and distal appendage recruitment

We next sought to determine the physiological role of myosin VI-OFD1 interaction by examining the effects of myosin VI depletion on centriole morphology and activity. To this end, we moved to retinal pigment epithelial (hTERT-RPE1) cells, diploid immortalised cells that maintain normal checkpoints on cell cycle progression and are commonly used for ciliogenesis assay. Initial characterisation of myosin VI knock-down (KD) RPE1 cells failed to detect major alterations in the structure of the centrioles, as evident by transmission electron microscopy (TEM) analysis (Supplementary Figure 3A). Both TEM and immunofluorescence analysis highlighted that the subcellular localisation of the centrosome, identified by pericentrin staining, is significantly altered upon myosin VI depletion, with an increased centrosome-plasma membrane distance in myosin VI KD cells (Supplementary Figure 3B).

Then, we focused on OFD1 behaviour at centrioles. OFD1 localisation is exquisitely limited to two specific pools - at the centrioles and at the centriolar satellites - that have different functions and regulations (Lopes et al. 2011; Tang et al. 2013; Wang et al. 2018). We first focused on the centriolar pool using the microtubules depolymerising drug nocodazole (Dammermann and Merdes 2002) to remove the satellites (Supplementary Figure 4A). To assess the amount of OFD1 at the centrioles, we calculated the fluorescence intensity at the centrioles’ spots, identified by the anti-centrin1 staining. To avoid biases due to the different phases of centriole duplication, we considered only cells with two centrioles, one of which marked by the distal appendage protein cep164 (Figure 2A). Our analysis revealed that myosin VI depletion causes an accumulation of OFD1 signal both in the mother centriole, identified with cep164, and in the daughter cep164-negative centriole (Figure 2B-C). Interestingly, the increase of OFD1 signal at the centrioles is not accompanied by a parallel increase of the protein level (Figure 2D), indicating a selective increased recruitment of OFD1 at the centrioles.

**Figure 2:**
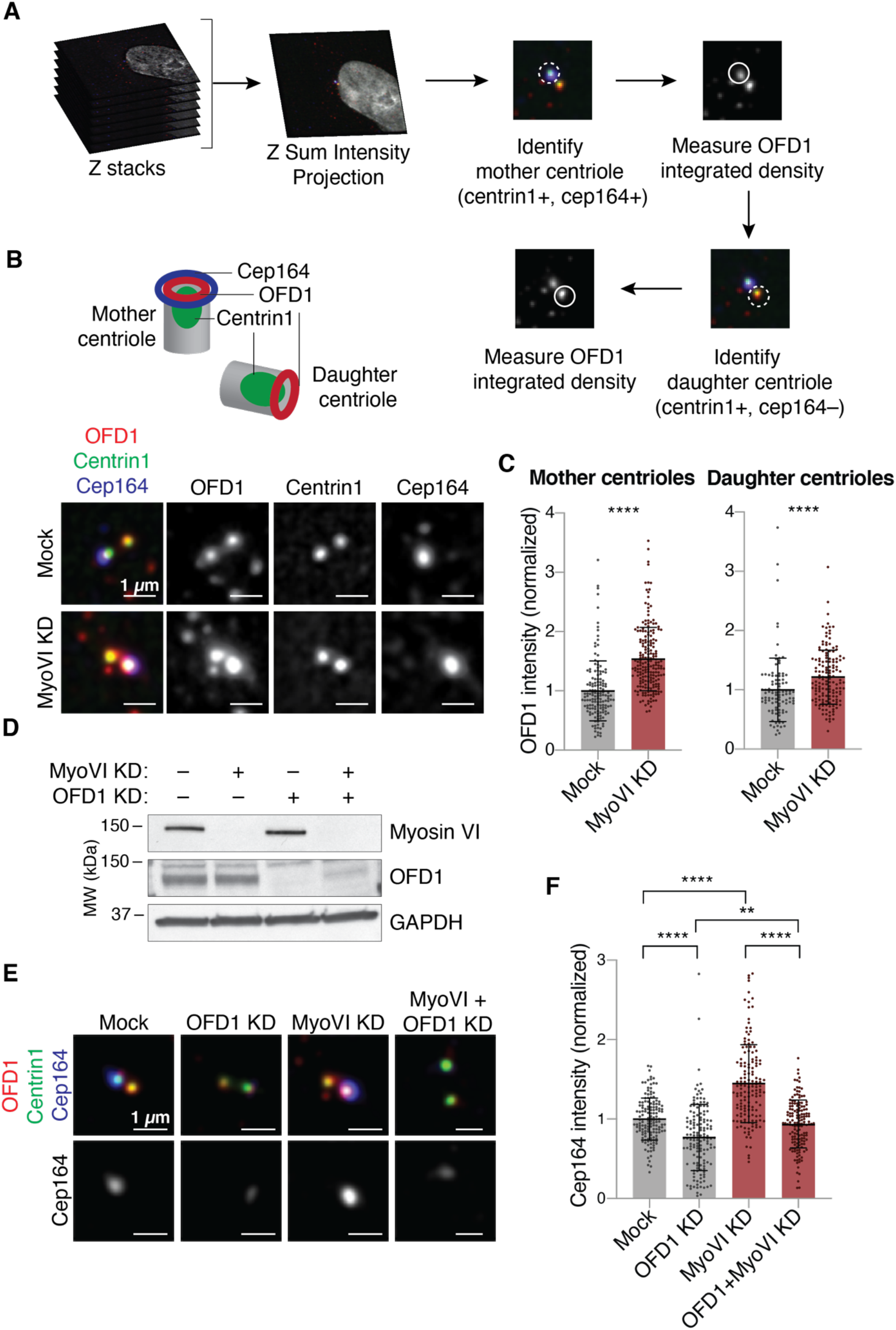
Myosin VI depletion leads to increased OFD1 and Cep164 recruitment at the centrioles. (A) A scheme of the IF analysis performed to calculate the total intensity of OFD1 staining at the mother and daughter centrioles. (B,C) IF analysis of centriole-associated OFD1 signal. hTERT-RPE1 cells were transfected with siRNA against myosin VI. Four days after transfection, cells were treated with nocodazole (1 hour, 6 µg/ml) and immunostained with anti-OFD1, anti-centrin1 and anti-cep164 antibodies. Mother centrioles were identified by the coincident staining of centrin1 and cep164, while daughter centrioles were centrin1-only stained. A scheme of the position of the markers used is depicted above. (B) Representative images, scale bar, 1 um. (C) Quantification of OFD1 intensity at the mother or daughter centrioles. Results are expressed as fold change with respect to mock average intensity. Bars represent mean ± SD. Mother centrioles: Mock, n=148 cells; MyoVI KD, n=199 cells, from four independent experiments. Daughter centrioles: Mock, n=101 cells; MyoVI KD, n=148 cells, from three independent experiments. **** P<0,0001 by Mann-Whitney test. (D) IB analysis of hTERT-RPE cells treated with the indicated siRNAs with anti-myosin VI and anti-OFD1 antibodies. Anti-GAPDH was used as loading control. (E-F) IF analysis of cep164 signal. hTERT-RPE1 cells were transfected with siRNA against myosin VI and/or OFD1. Four days after transfection, cells were treated with nocodazole (1 hour, 6 µg/ml) and immunostained with anti-OFD1, anti-centrin1 and anti-cep164 antibodies. (E) Representative images. Scale bar, 1 um. (F) Quantification of cep164 intensity at the mother centrioles. Results are expressed as fold change with respect to mock average intensity. Bars represent mean ±SD. Mock, n=147 cells; OFD1 KD, n=146 cells; MyoVI KD, n=150 cells; MyoVI + OFD1 KD, n=151 cells, from four independent experiments. ** P<0,005; **** P<0,0001 by Kruskal-Wallis test.

Previous studies demonstrated that OFD1 is required to constrain centriole elongation and to promote the recruitment of the distal appendages at the mother centriole (Wang et al. 2018; Singla et al. 2010). Confirming the aberrant accumulation of OFD1 at centrioles, we found that myosin VI depletion leads to an increased recruitment of the distal appendage protein cep164 at the mother centriole (Figure 2E-F). Moreover, the effect scored in myosin VI KD cells was rescued by the parallel depletion of OFD1, indicating that the cep164 accumulation is a secondary effect caused by OFD1 increase at centrioles (Figure 2E-F).

### Myosin VI depletion induces cell cycle arrest in non-tumoural cells through p53 activation

While exploring the effects of myosin VI KD, we unexpectedly observed a severe proliferation impairment in myosin VI-depleted RPE1 cells (Figure 3A). Similar results were obtained with additional siRNA oligos (Supplementary Figure 5A-C) and in BJ-hTERT cells, another non-tumoural cell line (Figure 3B). Analysis of the cell cycle by PI staining and FACS showed that myosin VI depletion induces an arrest in the G0/G1 phase, resulting in cellular senescence (Figure 3C-D). To investigate the cause of the cell cycle arrest, we performed immunoblot analysis and found that myosin VI-depleted cells display an increase in both p53 and p21 expression levels (Figure 3E). Consistently, double depletion of myosin VI and p53 by siRNA oligos rescues the proliferation of RPE1 cells (Figure 3F-G, Supplementary Figure 5A-C).

**Figure 3:**
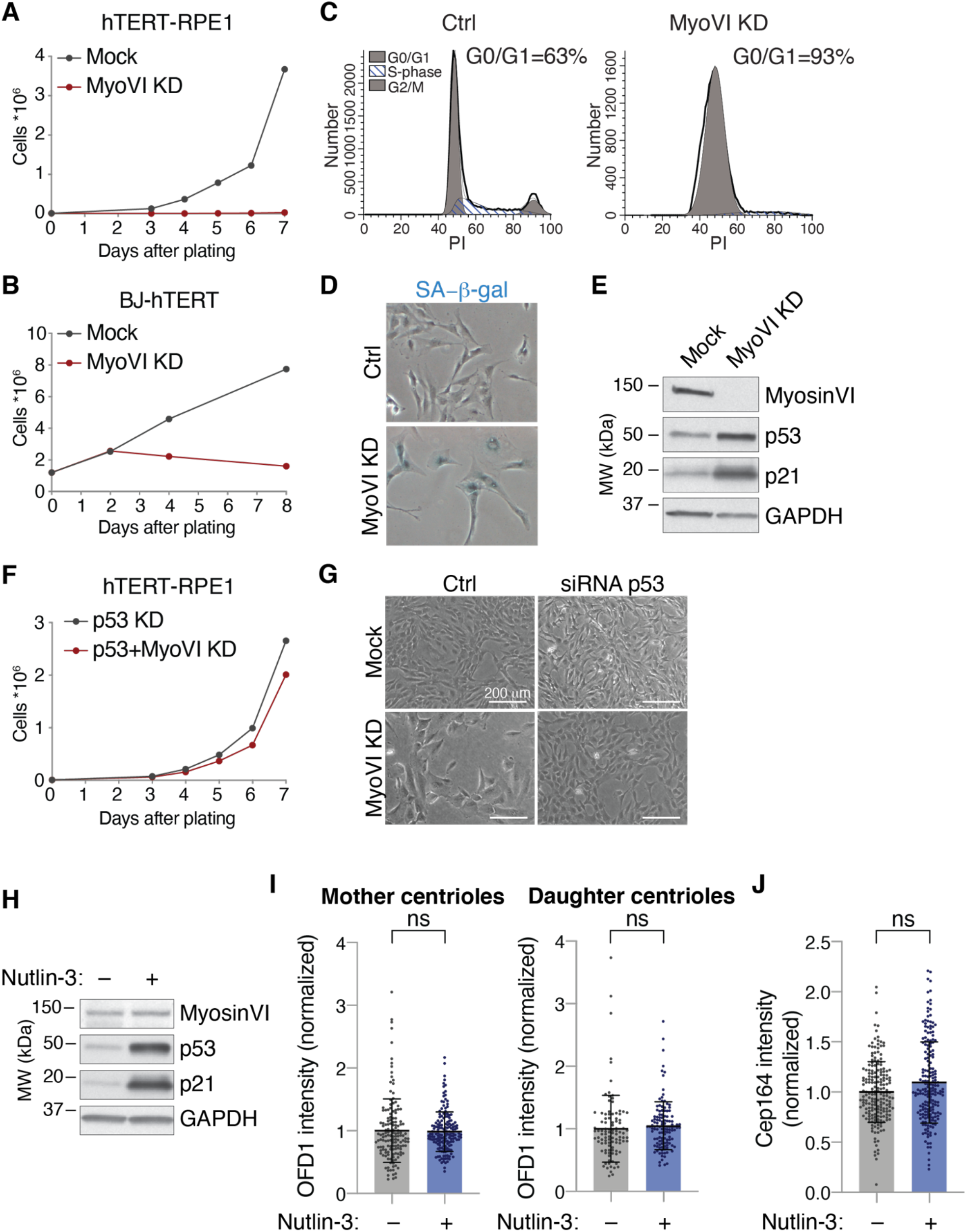
Myosin VI depletion leads to p53 activation and cell cycle arrest. (A) Growth curve of hTERT-RPE1 cells transfected with myosin VI siRNA. A representative plot of three independent experiments is shown. (B) Growth curve of BJ-hTERT cells transfected with myosin VI siRNA. A representative plot of two independent experiments is shown. (C) Analysis of DNA content in hTERT-RPE1 cells stably expressing a myosin VI shRNA. After 10 days of doxycycline induction, the cells were stained with Propidium Iodide (PI) and analysed by FACS. Ctrl = control cells, not induced. (D) Senescence-associated β-gal assay (SA-β-gal) of cells treated as in C. (E) IB analysis of hTERT-RPE1 cells transfected with myosin VI siRNA, with anti-myosin VI, anti-p53 and anti-p21 antibodies. Anti-GAPDH was used as loading control. (F) Growth curve of hTERT-RPE1 cells transfected with the indicated p53 and myosin VI siRNAs. A representative plot of three independent experiments is shown. (G) Representative bright field images of cells treated with the indicated p53 and myosin VI siRNAs. (H) IB analysis with anti-myosin VI and anti-OFD1 antibodies of hTERT-RPE cells treated with Nutlin-3, or not treated as control. Anti-GAPDH was used as loading control. (I) Quantification of OFD1 intensity at the mother or daughter centrioles. hTERT-RPE1 cells were transfected with siRNA against myosin VI. Four days after transfection, cells were treated with nocodazole (1 hour, 6 µg/ml) and immunostained with anti-OFD1, anti-centrin1 and anti-cep164 antibodies. Mother centrioles were identified by the coincident staining of centrin1 and cep164, while daughter centrioles were centrin1-only stained. Results are expressed as fold change with respect to mock average intensity. Bars represent mean ±SD. Mother centrioles: Mock, n=148 cells; MyoVI KD n=169 cells, from four independent experiments. Daughter centrioles: Mock n=101 cells; MyoVI KD; n=122 cells, from three independent experiments. ns, not significant by Mann-Whitney test. (J) Quantification of cep164 intensity at the mother centrioles in hTERT-RPE1 cells treated as in (I). Results are expressed as fold change with respect to mock average intensity. Bars represent mean ±SD. Mock, n=195 cells; MyoVI KD n=196 cells, from four independent experiments. ns, not significant by Mann-Whitney test.

To clarify if the increase in OFD1 and cep164 recruitment to the centrioles upon myosin VI depletion may be due to p53 accumulation and cellular senescence, cells were treated with Nutlin-3. This drug is a potent inhibitor of the p53 antagonist Mdm2 and induces p53 accumulation with consequent cell cycle arrest (Vassilev et al. 2004). After Nutlin-3 treatment, cells did not show any changes in the levels of OFD1 or cep164 at the centrioles (Figure 3H-J), confirming that the increased accumulation of OFD1 at centrioles we scored upon myosin VI depletion is specific to the lack of the myosin VI activity.

We then assessed if the activation of p53 upon myosin VI KD depends on defects occurring at centrioles. Indeed, alterations in the number or structure of the centrioles has been shown to cause cell cycle arrest, which is mediated by the activation of different pathways depending on the type of damage (Mikule et al. 2007; Ganem et al. 2014; Lambrus et al. 2016; Fong et al. 2016; Meitinger et al. 2016; Fava et al. 2017). To assess if the centrioles contribute to myosin VI depletion-induced p53 activation and cell cycle arrest, we sought to remove the centrioles from the cells and analyse cell proliferation. We used centrinone, a Plk4 inhibitor that blocks the formation of new centrioles (Wong et al. 2015), thus depleting them over a few cell cycles (Supplementary Figure 5D). To avoid the activation of p53 due essentially to centriole loss, we performed this experiment in 53BP1 KO RPE1 cells, in which p53 is not activated following centrinone treatment (Lambrus et al. 2016; Fong et al. 2016; Meitinger et al. 2016) (Supplementary Figure 5E). Strikingly, in centrinone-treated 53BP1 KO cells, myosin VI depletion caused p53 activation and cell cycle arrest similar to wild-type cells, as shown by Western blot and growth analysis (Supplementary Figure 5E, F).

Taken together, these results demonstrate that a p53-dependent cell cycle arrest occurs upon myosin VI depletion in non-tumoural cell lines, and that a general p53 activation does not induce OFD1 and Cep164 accumulation, as myosin VI KD does.

### Myosin VI controls the turnover of OFD1 at the centrioles

Given the increased recruitment of OFD1 at the centrioles upon myosin VI depletion and the localisation of the motor protein at the centrioles, we hypothesised that myosin VI could have a direct impact on OFD1 at the centrioles. Through Structured illumination microscopy (SIM) super-resolution microscopy, we confirmed the ring-like localisation of OFD1 at the distal tip of the mother and daughter centrioles in control cells (Wang et al. 2018). Conversely, we found that the lack of myosin VI caused an aberrant accumulation of OFD1 along the entire centriolar walls (Figure 4A).

**Figure 4:**
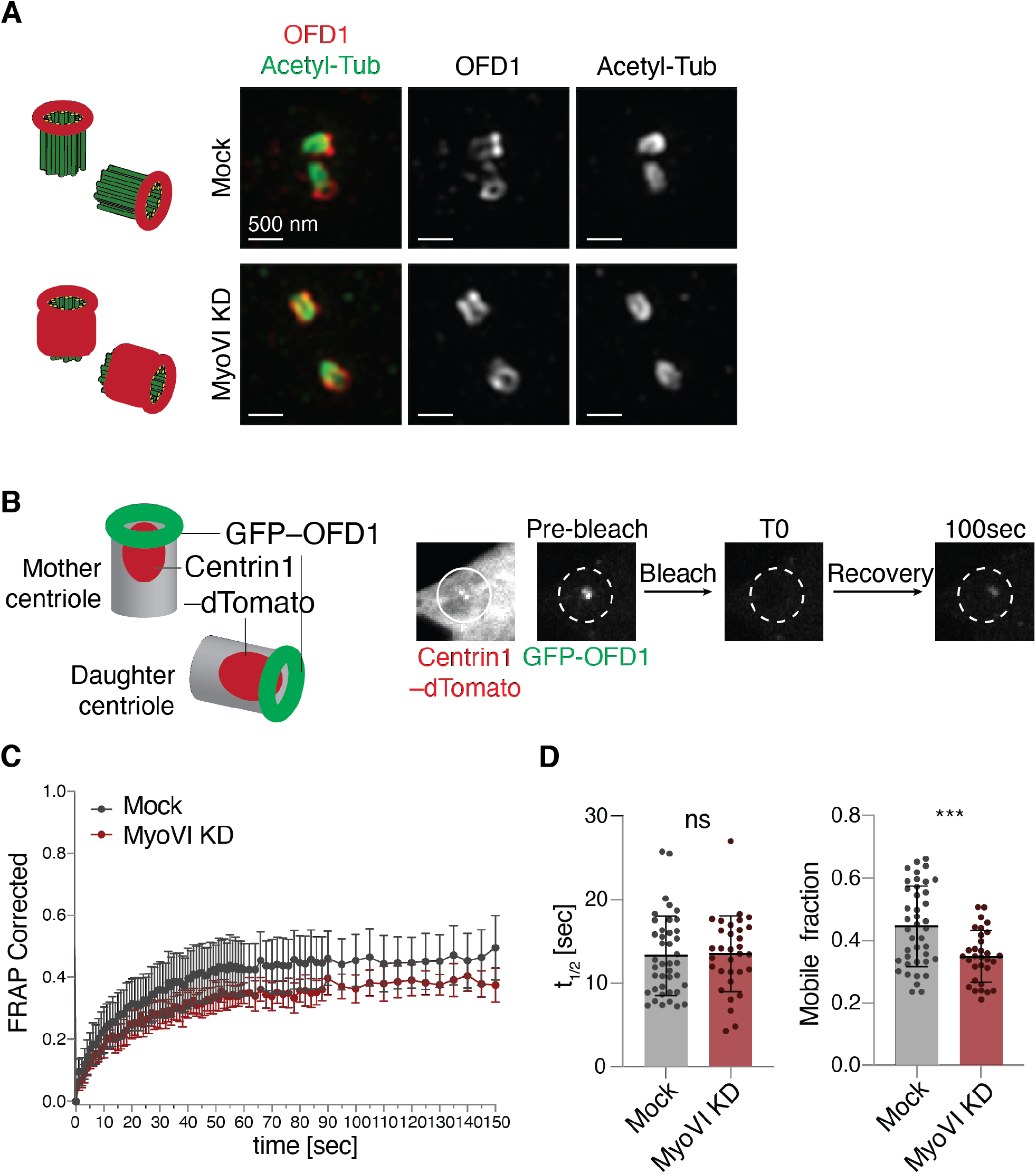
Lack of myosin VI alters the mobility and localisation of OFD1 at the centrioles. (A) Super-resolution analysis of OFD1 localisation at the centrioles. hTERT-RPE1 cells were transfected with siRNA against myosin VI, immunostained with anti-OFD1 and anti-acetylated tubulin antibodies and visualised using structured illumination microscopy (SIM). Representative images, scale bar, 500 nm. A scheme of the estimated localisation of OFD1 in the two conditions is depicted on the right side. (B-D) FRAP analysis of centriole-associated GFP-OFD1. hTERT-RPE cells stably expressing GFP-OFD1 and centrin1-dTomato were transfected with siRNA against myosin VI. After four days, cells were treated with nocodazole (1 hour, 6 µg/ml) and subjected to live-cell imaging. (B) Left: a scheme of the localisation of GFP-OFD1 and the centriole marker centrin1-dTomato. Right: a scheme of photobleaching and recovery of GFP-OFD1 at the centrioles. (C) A representative graph of one out of three experiments. For each time point, the fraction of recovery of GFP-OFD1 is shown. Results are expressed as means with 95% confidence interval. n=12 cells (Mock), n=13 cells (MyoVI KD). (D) Quantification of the half-time of fluorescence recovery (t_1/2_,) and of the mobile fraction of GFP-OFD1. Results are expressed as mean ±SD. n=41 cells (Mock), n=31 cells (MyoVI KD), from three independent experiments. ns, not significant; *** P<0,0005 by Unpaired T-test.

This result prompted us to analyse the behaviour of OFD1 by fluorescence recovery after photobleaching (FRAP) to determine the turnover of the protein. Centrioles were identified with centrin1-dTomato (Figure 4B), while the satellite pool of OFD1 was eliminated with nocodazole treatment during the live cell imaging (Supplementary Figure 4B). While the speed of recovery was not affected, myosin VI depletion caused a significant decrease in the mobile fraction of OFD1, indicating the presence of a stable pool of OFD1 at the centrioles that cannot be mobilised (Figure 4C, D).

Collectively, these data indicate that in the absence of myosin VI, OFD1 cannot be removed from the centrioles and thus aberrantly accumulates at the centrioles.

### Myosin VI removes OFD1 from the satellites

Besides the centrioles, OFD1 also localises at the centriolar satellites in cycling cells. Previous studies showed that serum starvation induces OFD1 removal from the satellites by autophagy, a process that is required for primary ciliogenesis (Tang et al. 2013; Morleo et al. 2020). Interestingly, myosin VI is a known regulator of autophagy, being essential for autophagosome-lysosome fusion during autophagosome maturation (Tumbarello et al. 2012). To determine if myosin VI exerts a role on satellites, we first examined the effect of myosin VI depletion using the satellite marker PCM1, which is virtually absent at the centrioles (Dammermann and Merdes 2002). Although we could observe a strong reduction of PCM1 intensity around centrioles in myosin VI KD cells (Figure 5A and B), cells treated with Nutlin-3 showed a similar decrease (Figure 5C), suggesting that the p53 activation induced by myosin VI KD causes satellites loss. Consistently, myosin VI depletion does not affect PCM1 intensity in RPE1 p53 KO cells (Figure 5D).

**Figure 5:**
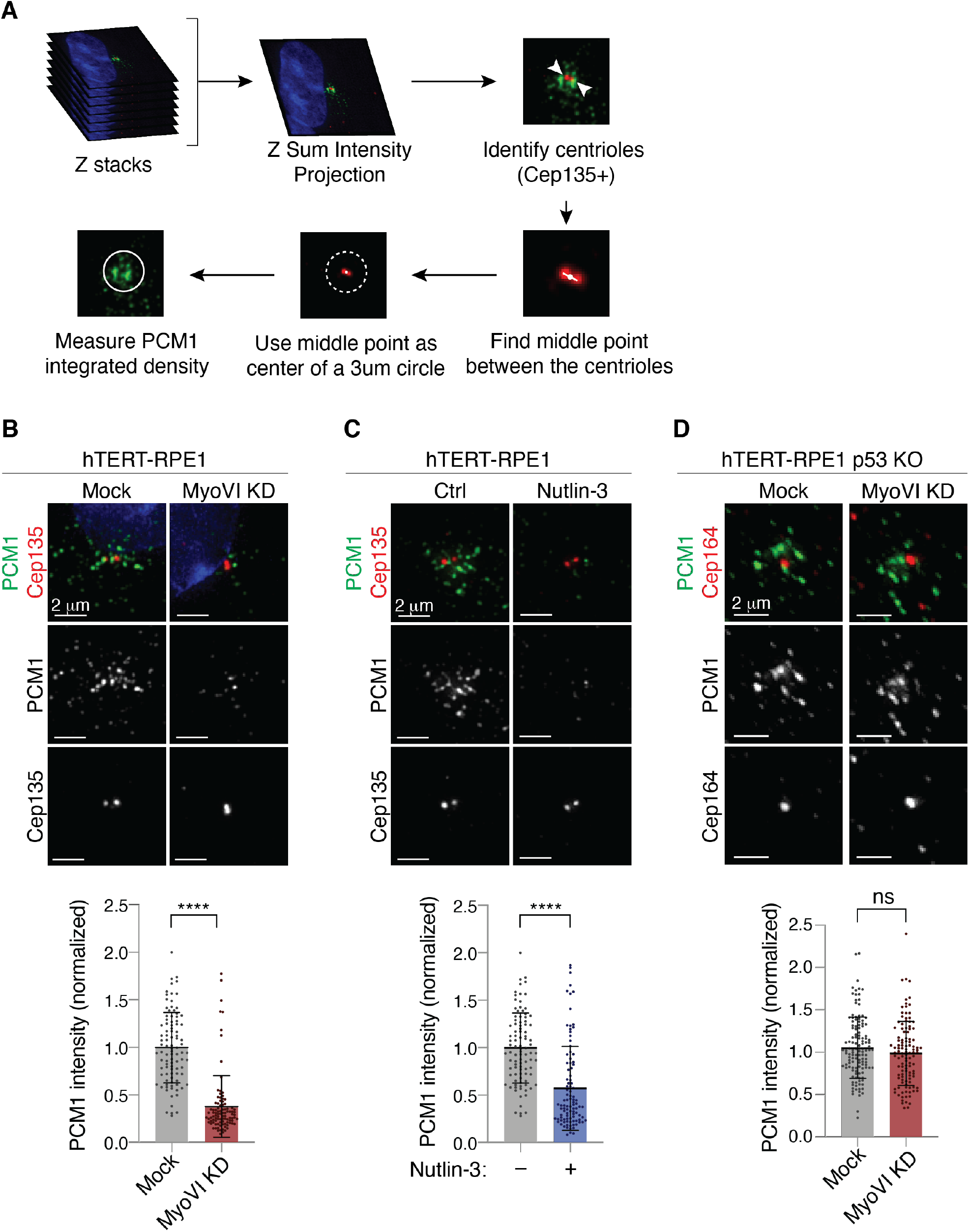
Myosin VI depletion affects the centriolar satellites via p53. (A) A scheme of the IF analysis performed to calculate the total intensity of satellite staining that surrounds the centrioles. The centriole marker cep135 or cep164 are used to define the centre of a 3 µm circle, in which the intensity of the satellite marker PCM1 was calculated. (B) IF analysis of PCM1 signal. hTERT-RPE1 cells were transfected with siRNA against myosin VI and immunostained with anti-PCM1 and anti-cep135 antibodies. Upper panel, representative images, scale bar, 2 µm. Lower panel, quantification of PCM1 intensity. Results are expressed as fold change with respect to mock average intensity. Bars represent mean ±SD. Mock, n=96 cell; MyoVI KD, n=98cells, from two independent experiments. **** P<0,0001 by Mann-Whitney test. (C) IF analysis of PCM1 signal. hTERT-RPE1 cells were treated with Nutlin-3 for 24 hours and immunostained with anti-PCM1 and anti-cep135 antibodies. Panels as in B. Mock, n=96 cells; Nutlin-3, n=100 cells, from two independent experiments. **** P<0,0001 by Mann-Whitney test. (D) IF analysis of PCM1 signal. hTERT-RPE1 p53 KO cells were transfected with siRNA against myosin VI and immunostained with anti-PCM1 and anti-cep164 antibodies. Panels as in B. Quantification of PCM1 intensity refers to a 3 µm circle around the mother centriole, identified with anti-cep164 staining. Mock, n=128 cells; MyoVI KD, n=114 cells from three independent experiments. ns, not significant by Mann-Whitney test.

Based on these results, we used RPE1 p53 KO cells to analyse the effect of myosin VI depletion on the satellite pool of OFD1. To eliminate the contribution of the centriolar pool of OFD1, we identified the satellites via PCM1 staining and calculated OFD1 intensity only in the area covered by PCM1 (Figure 6A). As previously shown (Tang et al. 2013; Akhshi and Trimble 2021), serum starvation induces the removal of OFD1 from the satellites. We could observe that this event is significantly impaired by the parallel depletion of myosin VI (Figure 6B). While serum starvation induces a decrease in OFD1 total protein levels (Akhshi and Trimble 2021; Morleo et al. 2020), the same effect is weaker in myosin VI-depleted cells (Figure 6C).

**Figure 6:**
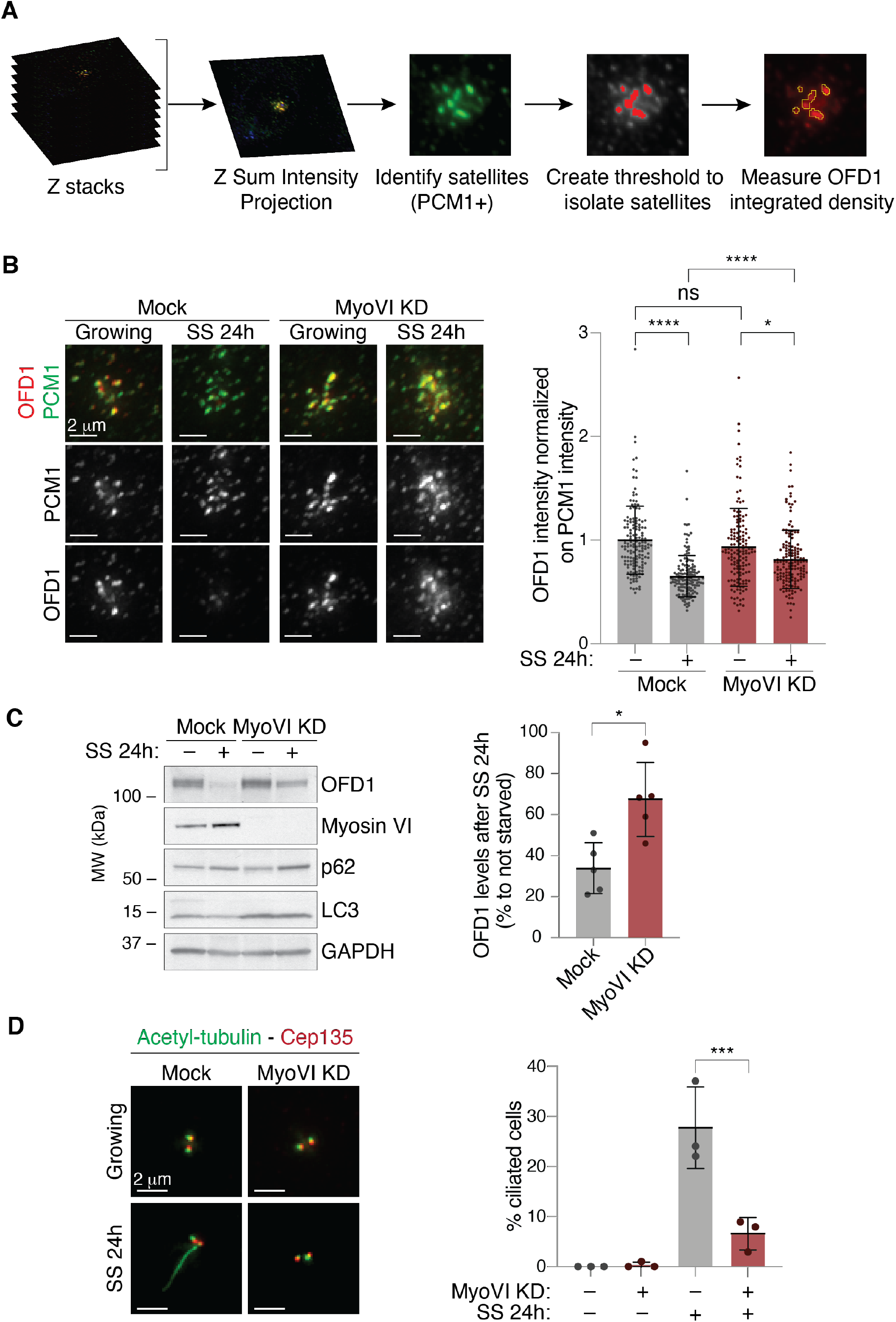
Myosin VI contributes to OFD1 removal from the centriolar satellites required for ciliogenesis. (A) A scheme of the IF analysis performed to calculate the total intensity of OFD1 staining at the centriolar satellites. (B) IF analysis of OFD1 signal at the centriolar satellites upon serum starvation. hTERT-RPE1 p53 KO cells were transfected with siRNA against myosin VI. After four days, cells were fixed (growing) or serum starved for 24 hours (SS). Cells were immunostained with anti-OFD1 and anti-PCM1 antibodies. The intensity of OFD1 signal in the area covered by PCM1 was quantified and normalised against the intensity of PCM1 staining in the same area. Left, representative images, scale bar, 2 µm. Right, results are expressed as fold change with respect to mock average intensity. Bars represent mean ±SD. Mock_growing, n=150 cells; Mock_SS, n=149 cells; MyoVI KD_growing, n=149 cells; MyoVI KD_SS, n=150 cells, from three independent experiments. ns, not significant; * P<0,05; **** P<0,0001 by Kruskal-Wallis test. (C) IB analysis of OFD1 after serum starvation in control and myosin VI-depleted cells. hTERT-RPE1 p53 KO cells were transfected with siRNA against myosin VI. After four days, cells were serum starved for 24 hours (SS). Lysates were analysed by IB with anti-OFD1, anti-myosin VI, anti-p62, and anti-LC3 antibodies. Anti-GAPDH was used as loading control. The amount of OFD1 protein was normalised against GAPDH signal and is expressed as percentage of OFD levels in cells grown in serum-starved conditions compared to cells grown in media containing serum. Bars represent mean ±SD. n=5 independent experiments. * P<0,05 by Kruskal-Wallis test. (D) IF analysis of primary cilium upon serum starvation. hTERT-RPE1 p53 KO cells were transfected with siRNA against myosin VI. After four days, cells were fixed (growing) or serum starved for 24 hours (SS). Cells were immunostained with anti-acetylated tubulin (to identify the cilia), anti-cep135 (to identify the centrioles) and DAPI. Left, representative images, scale bar, 2 µm. Right, results are expressed as fold change with respect to mock average intensity. Bars represent mean ±SD. n=3 independent experiments. 100-200 cells/condition were counted for each experiment.

Ciliogenesis in tissue culture is mainly initiated by serum starvation, which arrests the cell cycle and triggers autophagy (Tang et al. 2013; Morleo et al. 2020). Our data support the idea that myosin VI contributes to OFD1 degradation induced by serum starvation. Thus, we examined the formation of primary cilia in myosin VI-depleted cells. To avoid confounding effect due to the cell cycle arrest induced by myosin VI depletion, we performed the experiment in RPE p53 KO cells. Remarkably, while around 30% of the p53 KO cells were found ciliated after starvation even in the absence of G1 arrest, myosin VI-depleted cells completely failed to form cilia (Figure 6D). These data imply a requirement of myosin VI activity for primary cilium formation.

## Discussion

Here, we identified and characterised a novel myosin VI interactor, OFD1, whose turnover is modulated by myosin VI both at the centrioles and at the centriolar satellites. Myosin VI activity on OFD1 appears to be critical for maintaining the correct amount of the distal appendages at the mother centriole and for the removal of OFD1 from the centriolar satellites to promote primary ciliogenesis. Thus, OFD1 can be added to the list of specialised cargo-adaptor proteins that link myosin VI to distinct cellular compartments and processes (Magistrati and Polo 2020). Importantly, both myosin VI short and long isoforms (Wollscheid et al. 2016) interact with OFD1, indicating that this functional interaction is conserved in all cells and is maintained upon epithelial cell polarisation where a switch between isoforms was observed (Biancospino et al. 2019; Buss et al. 2001)

At the centrioles, OFD1 localisation is restricted to the distal tip (Figure 4A) (Tang et al. 2013). Our FRAP analysis determined that the 45% of OFD1 present at the centrioles is mobile and rapidly exchanges with the cytoplasmic pool. In the absence of myosin VI, this fraction is reduced, allowing accumulation of OFD1 on the entire microtubule wall. At present, the mechanism by which myosin VI controls OFD1 turnover remains unclear. Since the speed of recovery is not affected (t1/2 in FRAP analysis, Figure 4D), it is conceivable that myosin VI is needed for short-range transport and may exploit the actin filaments that are nucleated by the centrosome to promote the turnover of OFD1 via its motor activity. In this context, it is noteworthy that the SAH domain involved in myosin VI dimerisation and required for the movement (Mukherjea et al. 2014) is critical for the interaction between the two proteins (Figure 1E). At the distal tip, OFD1 could be stabilised by other proteins like C2CD3 (Thauvin-Robinet et al. 2014) that protect this centriolar protein from myosin VI-mediated removal.

Recent studies have highlighted a synergistic interplay between actin and microtubule dynamics in several cellular compartments. Actin-microtubule crosstalk is functionally relevant for mitotic spindle positioning (Farina et al. 2019; Inoue et al. 2019) or for cell shape and polarity during cell migration, and many proteins that mediate actin–microtubule interactions have already been identified (Dogterom and Koenderink 2019). Our data raise the possibility that myosin VI is one of these critical modulators at the centrosomes where it could exploit the actin-based network surrounding the centrosomes and the microtubule interactors that we have identified with the proteomic approach (i.e. Numa, Supplementary Table 1). Further investigations are needed to uncover the functional implications of this intriguing hypothesis that is supported by the recent identification of a Gαi-LGN-NuMA-dynein axis activated upon Shh-Smo induction to promote ciliogenesis (Akhshi and Trimble 2021), and which is consistent with the aberrant positioning of the centrosomes observed in the absence of myosin VI (Supplementary Fig. 3B).

Our study also unveils an unprecedent phenotype of cell-cycle arrest and senescence following myosin VI depletion in p53-proficient cells. This effect was not evident in cancer cell lines, suggesting that myosin VI contributes to a pathway that sustains proliferation and maintains the cell cycle in check, and that becomes deregulated during carcinogenesis. While this possibility is currently under investigation, the cause of this phenotype is certainly unrelated to centriolar biology as demonstrated by the centrinone experiment (Supplementary Figure 5B,C). Another surprising finding of our study is that p53 activation is responsible for the satellite dispersion occurring upon myosin VI depletion, which was established by Nutlin-3 treatment that phenocopies myosin VI depletion, as well as by the rescue of the phenotype obtained in p53 KO cells. Thus, our study highlights a new intriguing function of p53 at satellites. Determination of how p53 senses the lack of myosin VI is an important area of future work that is predicted to shed light also on this phenotype.

Finally, our study identified for the first time myosin VI as a critical regulator of ciliogenesis. Based on its role in other cellular compartments, it is possible that myosin VI may act as an actin anchor for the basal body, by docking the mother centriole at the plasma membrane, or as an actin-based motor contributing to the trafficking towards the ciliary pocket (Akhshi and Trimble 2021). Nonetheless, our evidence suggests that myosin VI depletion affects autophagy-mediated removal of OFD1 from the satellites, a prerequisite for ciliogenesis (Akhshi and Trimble 2021; Morleo et al. 2020; Tang et al. 2013). Several findings link myosin VI to autophagy and autophagy receptors (Magistrati and Polo 2020; Tumbarello et al. 2012), and emerging evidence suggests that autophagy and ciliogenesis influence each other (Morleo and Franco 2019; Pampliega and Cuervo 2016). Mechanistically, myosin VI has been shown to act at the damaged mitochondria where it mediates their engulfment and clearance via the formation of F-actin cages (Kruppa and Buss 2018). The interaction with OFD1 suggests that myosin VI may be directly involved in the first phases of the autophagosome formation, although implication of F-actin cages cannot be excluded. Whatever the case, the identification of myosin VI as OFD1 regulator has important implications for a variety of ciliopathy syndromes in which OFD1 mutations compromise ciliogenesis (Singla et al. 2010).

## Data availability

The mass spectrometry raw datasets were deposited in PRIDE database and can be accessed through ProteomeXchange with the following Project Name: Pulldown CG42797 vs S2 cell lysate and Project accession: PXD026697. Full list of the specific interactors is provided in Supplementary Table 1.

Acknowledgements

We thank Brunella Franco for DNA constructs, Francesca Farina and Luca Fava for the generation of cell lines. We thank Paolo Soffientini, Angela Cattaneo, Massimiliano Garrè and Emanuele Martini at Cogentech facilities (Milan, Italy) for support in mass spectrometry and FRAP analysis. We are grateful to Wessen Maruwge for English language editing. This work was supported by the Associazione Italiana per Ricerca sul Cancro, (Investigator grant 2017-19875 to S.P.). Elisa Magistrati’s work is supported by the Associazione Italiana per la Ricerca sul Cancro. E.Mag. was and G.S. is a PhD student at the European School of Molecular Medicine (SEMM).

## Author Contributions

Conceptualisation: E.Mag, and S.P; E.Mag, performed all the experiments with the help of G.M. and E.Mas.; M.L.F. aided in the imaging acquisition and SIM experiments; G.B. and A.M performed the EM experiment; M.B.D contributed to the planning and interpretation of data; S.P. coordinated the team, designed and supervised the project; E.Mag. and S.P. wrote the paper with contributions from all authors.

---

**Supplementary Figure 1:**
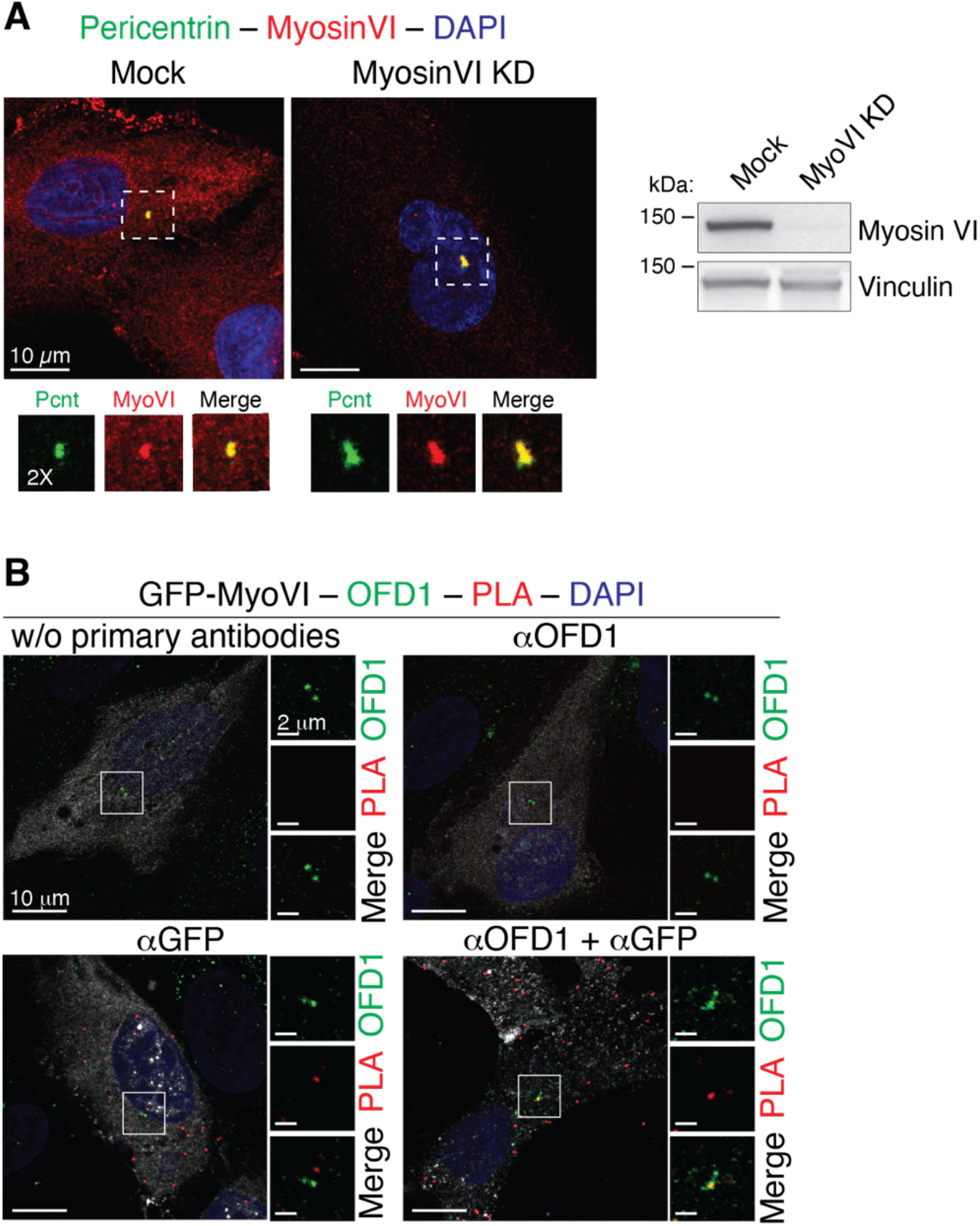
Myosin VI localisation at the centrosome. (A) hTERT-RPE1 cells were transfected with siRNA against myosin VI and immunostained with anti-myosin VI and anti-pericentrin antibodies. Co-localisation between myosin VI and pericentrin, maintained also in myosin VI KD cells, does not allow us to draw definitive conclusions about myosin VI centrosome localisation. Representative images are shown, scale bar, 10 µm. (B) Proximity ligation assay (PLA) in hTERT-RPE1 cells overexpressing GFP-myosin VI_short_ full-length, using anti-OFD1 and anti-GFP antibodies. After PLA assay, counterstaining with fluorescently-labelled secondary antibodies allowed the detection of OFD1 and GFP localisation. Representative images of technical controls with single or no primary antibodies are shown. Scale bar, 10 µm (2 µm for the magnification).

**Supplementary Figure 2:**
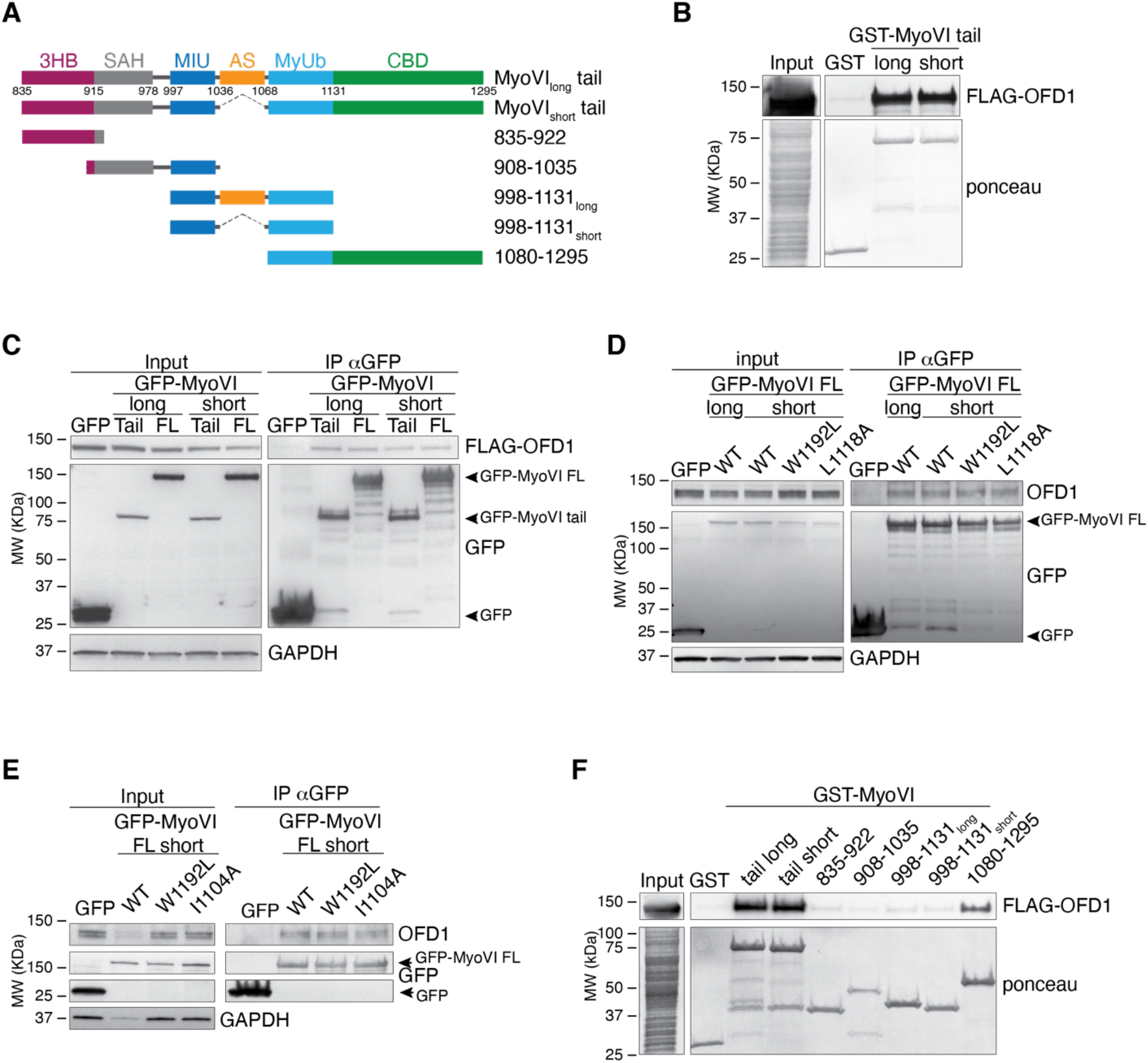
Characterization of the myosin VI minimal region of binding to OFD1. (A) A scheme of the structure and domain organisation of myosin VI. The tail domain is composed of a three-helix bundle (3HB), a single alpha helix (SAH), two ubiquitin binding regions (MIU = motif interacting with ubiquitin; MyUb = Myosin VI ubiquitin binding domain), and a cargo binding domain (CBD). Between the MIU and the MyUb, an alternative spliced region (AS, in orange) is present in myosin VI_long_ isoform, while it is absent in the myosin VI_short_ isoform. (B) GST pulldown assay using the indicated myosin VI tail (835-1295) constructs or GST alone as control and lysates from HEK293T cells transfected with Flag-OFD1 construct. IB was performed with anti-Flag antibody. Ponceau staining as indicated. (C) Total lysates from HEK293T transfected with Flag-OFD1 and the indicated GFP-myosin VI constructs (or GFP as control) were IP with anti-GFP antibody-conjugated beads. IB was performed with anti-OFD1 and anti-GFP antibodies. Anti-GAPDH was used as loading control. FL = full length. (D-E) Total lysates from HEK293T transfected with GFP (as control) or GFP-myosin VI constructs containing the indicated single-point mutations were IP with anti-GFP antibody-conjugated beads. IB was performed with anti-OFD1 and anti-GFP antibodies. Anti-GAPDH was used as loading control. FL = full length. (F) GST pulldown assay using the indicated myosin VI constructs or GST alone as control and lysates from HEK293T cells transfected with Flag-OFD1 construct. IB was performed with anti-Flag antibody. Ponceau staining as indicated.

**Supplementary Figure 3:**
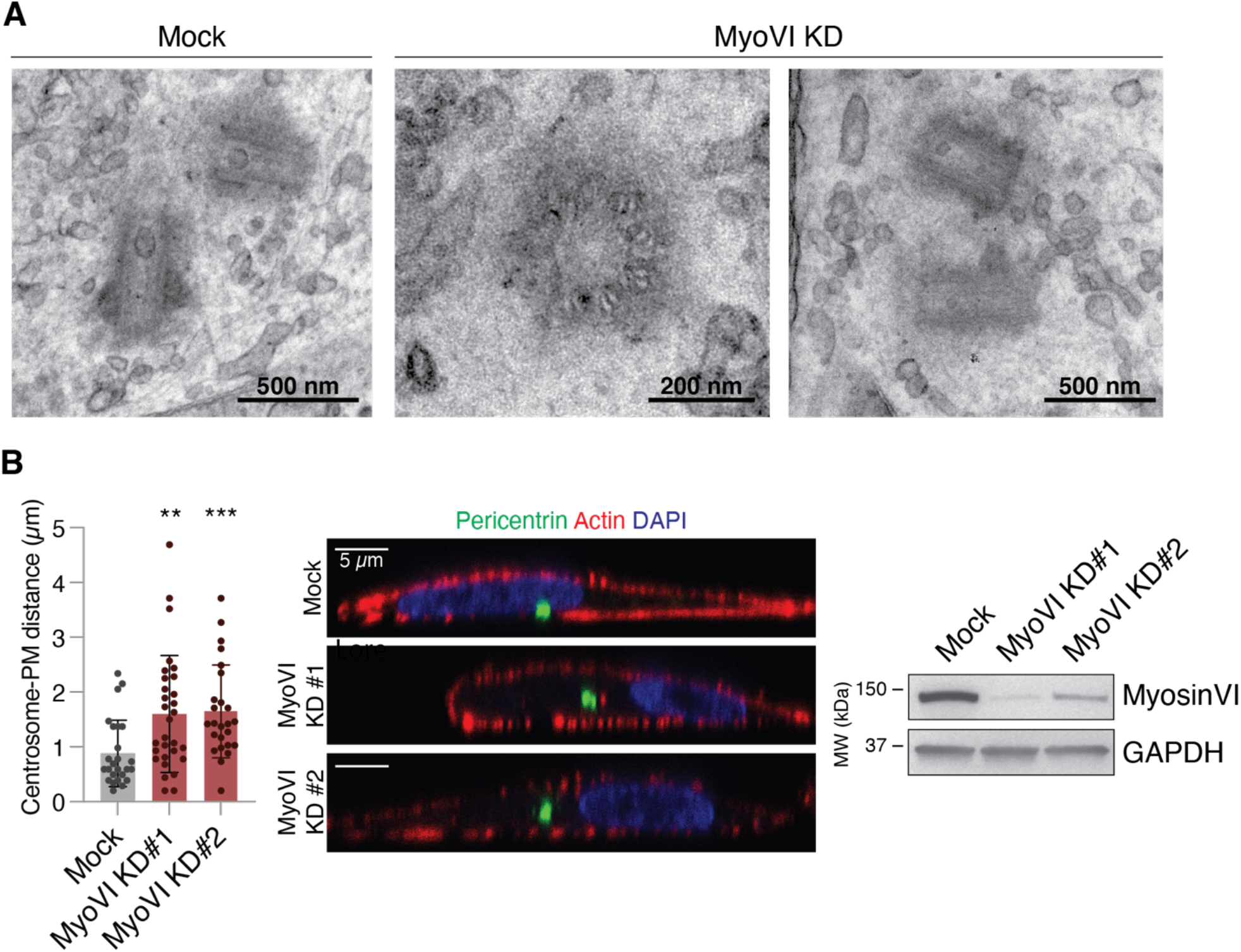
Myosin VI depletion does not affect centrioles ultrastructure. (A) Transmitted electron microscopy (TEM) analysis of centriole structure in hTERT-RPE1 control and myosin VI-depleted cells. Representative images are shown, scale bar as indicated. (B) Depletion of myosin VI leads to displacement of the centrosomes from the cell cortex. hTERT-RPE1 cells were transfected with the indicated siRNA oligos and, after 96 hours, were stained with anti-pericentrin to mark centrioles, phalloidin-TRITC to mark cortical actin, and DAPI to mark the nuclei. Left, the distance of the centrosome from the plasma membrane was calculated using ImageJ from XZ-axis images of the cells. The mean distance ± SD is reported. n=2 independent experiments. N=25-30 cells/condition were counted. ** P<0,005; *** P<0,001 by Kruskal-Wallis test. Centre, representative images are shown; scale bar, 5 µm. Increased distance between centrioles and nuclear membrane is evident. Right, IB analysis showing myosin VI depletion.

**Supplementary Figure 4:**
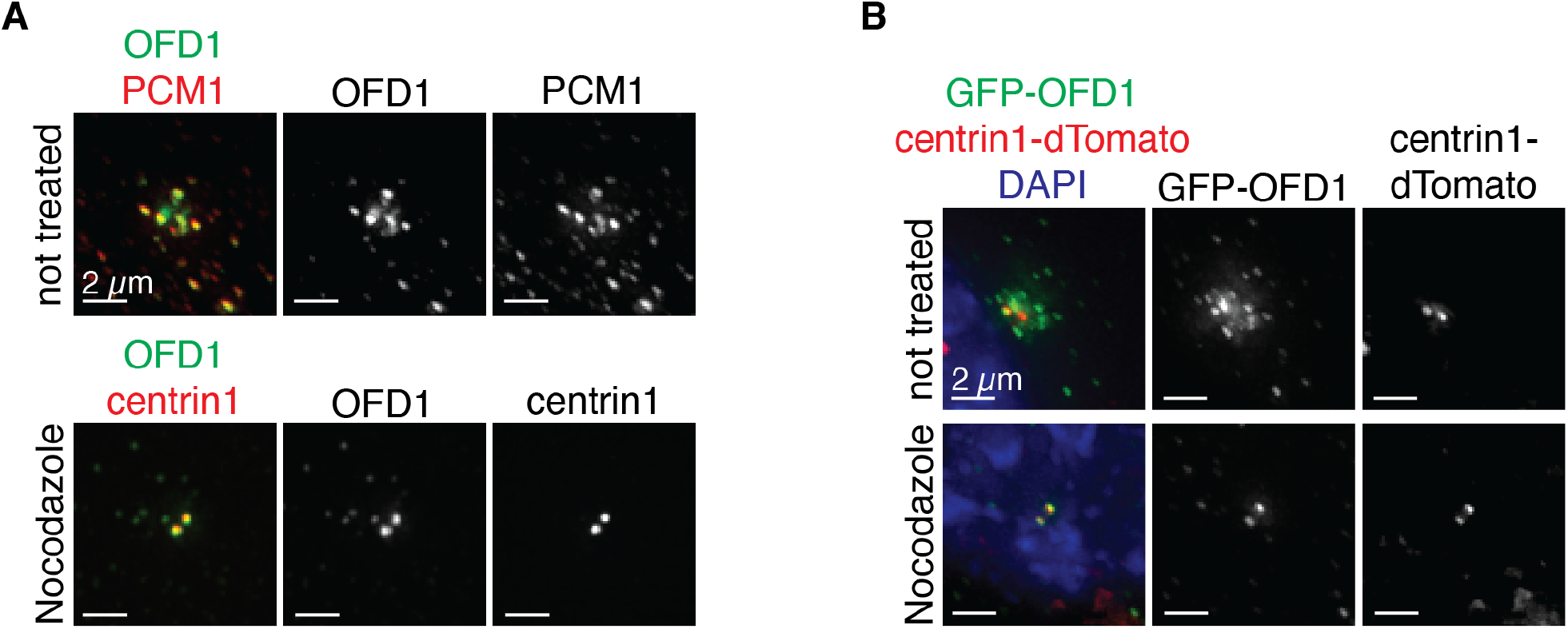
Nocodazole treatment leads to the dispersion of OFD1 satellite pool. (A) hTERT-RPE1 cells were treated with nocodazole at 6 µg/ml for 1 hour prior fixation to scatter the centriolar satellites and remove OFD1 satellites pool. Cells were immunostained with the anti-OFD1 antibody and the indicated markers for centriolar satellites (anti-PCM1) and centrioles (anti-centrin1). Representative images are shown; scale bar, 2 µm. (B) Effect of nocodazole treatment on GFP-OFD1 localisation in hTERT-RPE cells stably expressing GFP-OFD1 and centrin1-dTomato, used in FRAP experiments shown in Figure 4. Representative images are shown; scale bar, 2 µm.

**Supplementary Figure 5:**
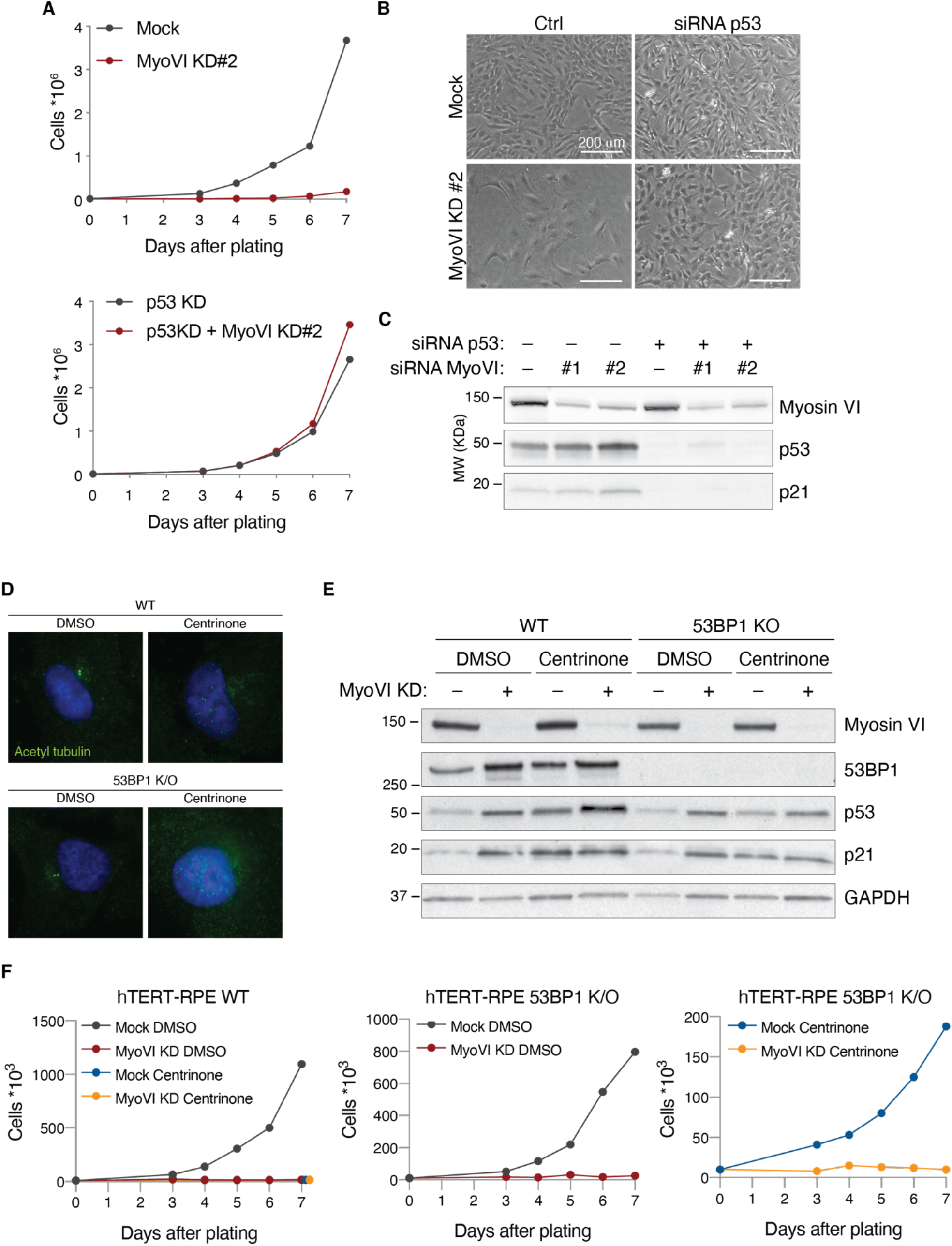
p53 activation in myosin VI-depleted cells is not due to its effect on centrioles. (A-C) Severe proliferation impairment in myosin VI-depleted RPE1 cells is reproducible using a second siRNA oligo against myosin VI (KD#2). (A) Growth curve of hTERT-RPE1 cells transfected with a second myosin VI siRNA oligo +/-p53 siRNA oligo. A representative plot of three independent experiments is shown. (B) Representative bright field images of cells treated with the indicated p53 and myosin VI siRNAs. (C) IB analysis of control and myosin VI-depleted hTERT-RPE cells with anti-myosin VI, anti-p53 and anti-p21 antibodies. Anti-GAPDH was used as loading control. (D) hTERT-RPE1 wild-type or 53BP1 KO cells were treated with centrinone or DMSO (as control) and immunostained with anti-acetylated tubulin antibody and DAPI to verify the presence or absence of centrioles. (E) Analysis of hTERT-RPE wild-type or 53BP1 KO cells transfected with myosin VI siRNA and treated with centrinone or DMSO (as control) by IB, with anti-myosin VI, anti-53BP1, anti-p53 and anti-p21 antibodies. Anti-GAPDH was used as loading control. (F) Growth curves of hTERT-RPE1 wild-type or 53BP1 KO cells transfected with myosin VI siRNA and treated with centrinone or DMSO. Representative plots of three independent experiments are shown.

**Supplementary Table 1.**
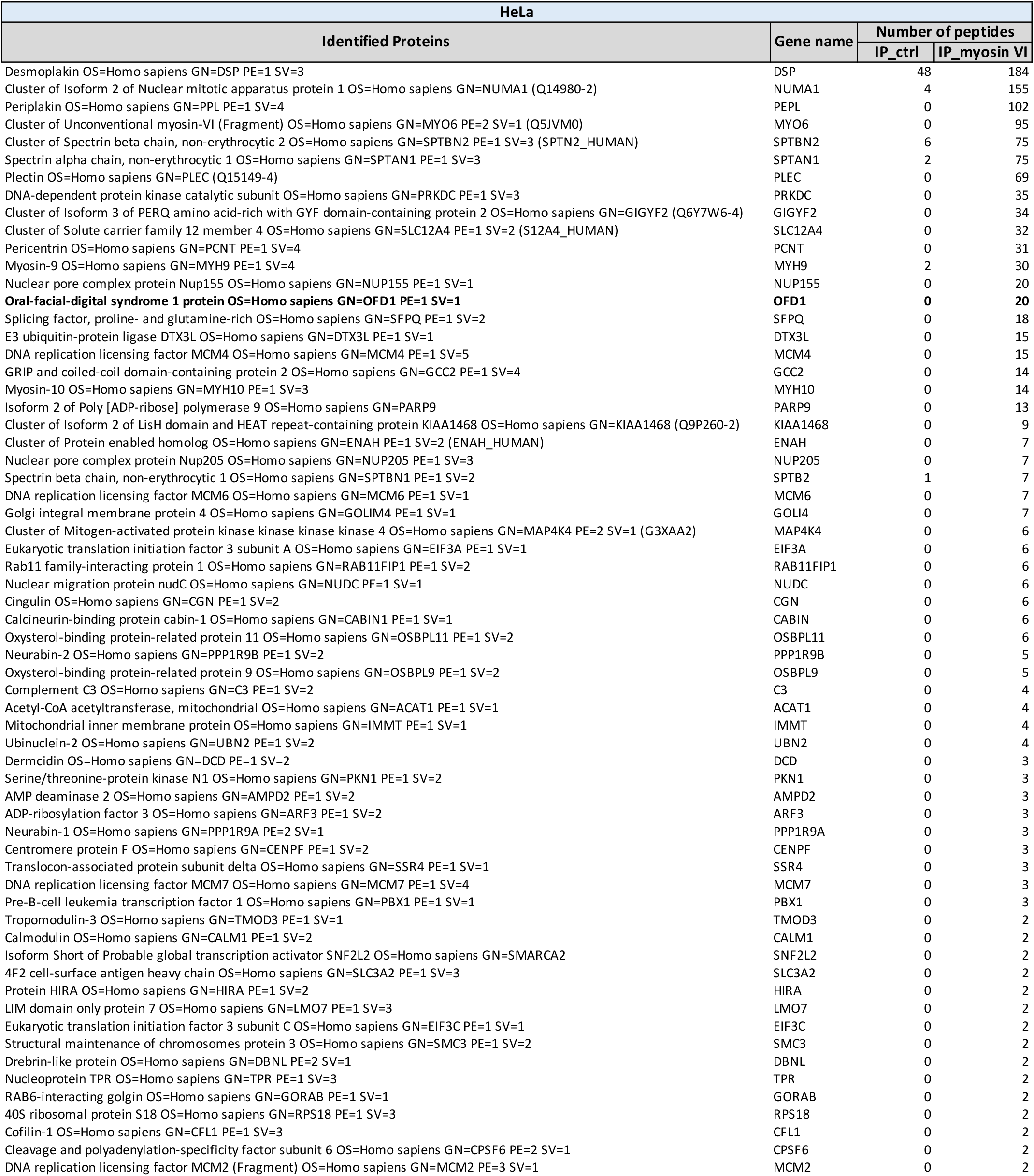
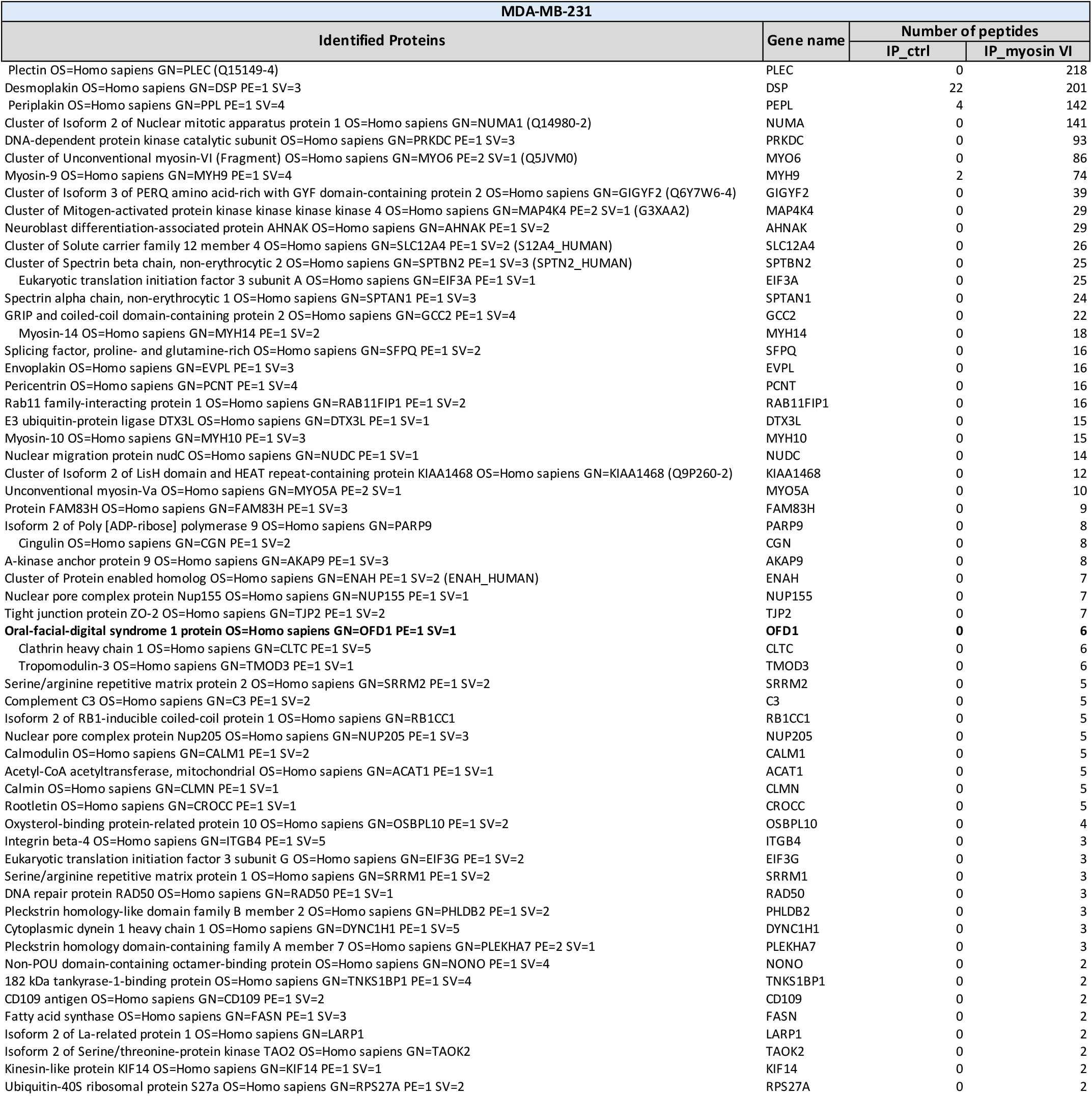
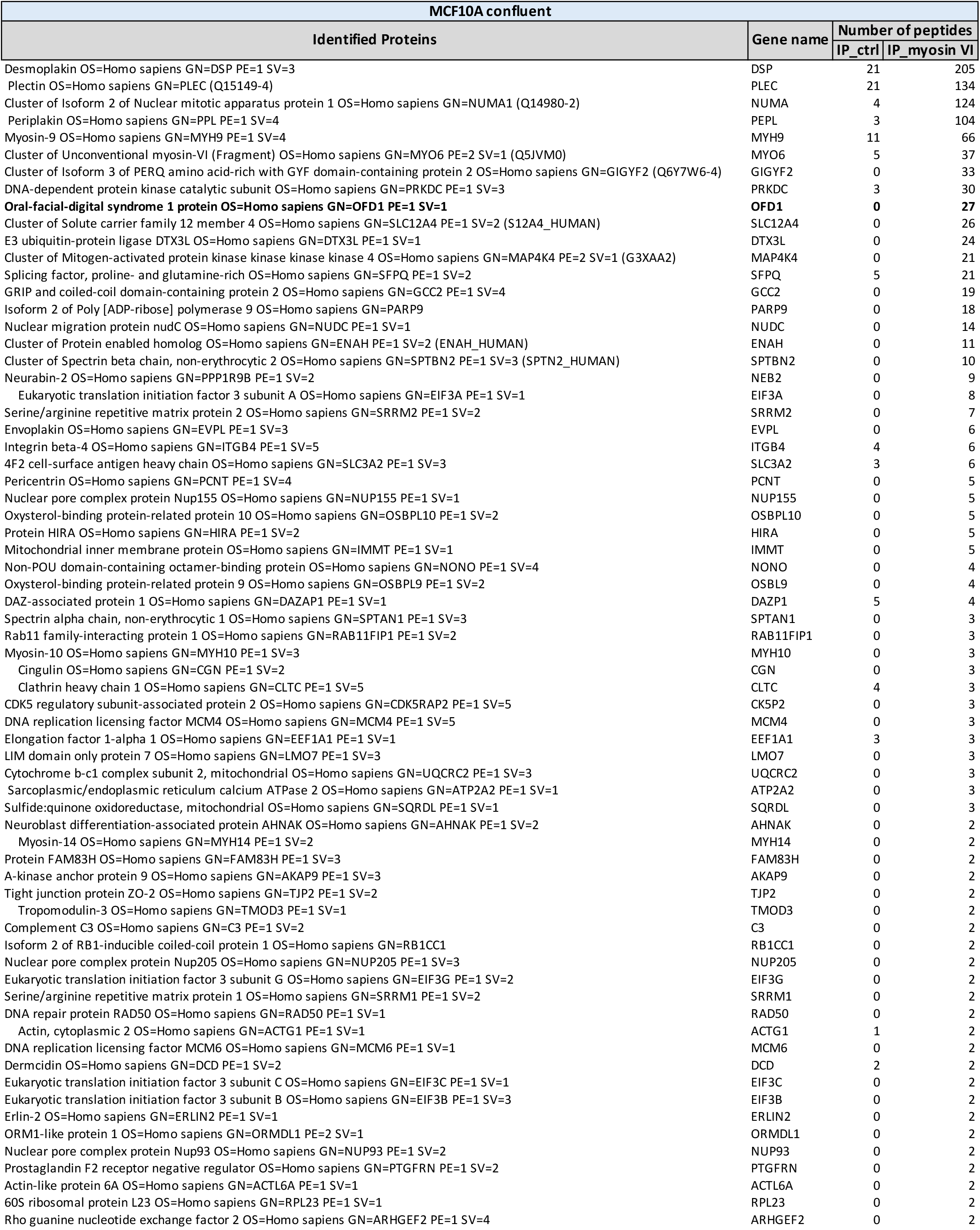
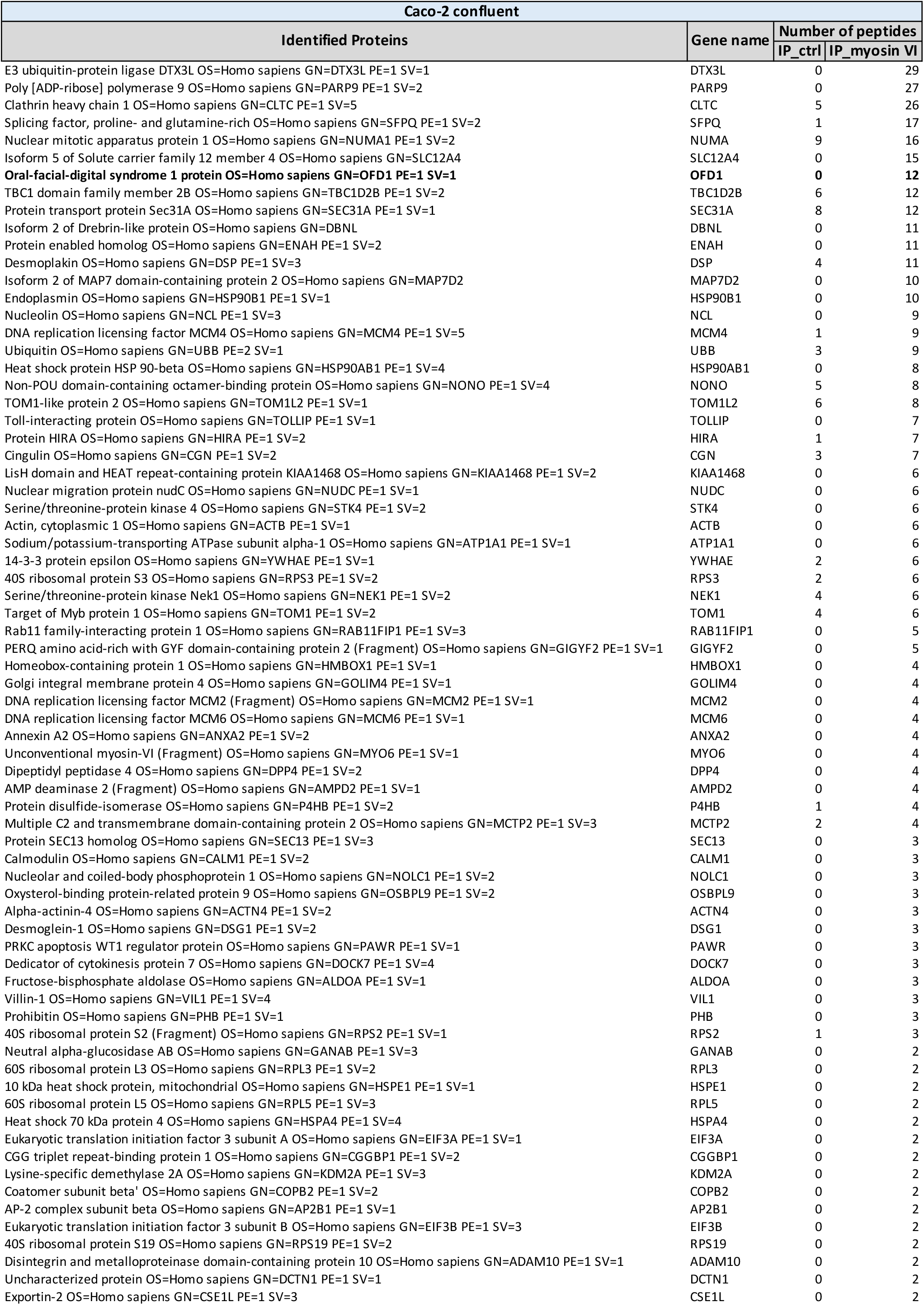

## Methods

### Constructs

pGEX-myosin VI tail (835-1295) and pEGFP-myosin VI full-length (FL) constructs were previously described (Wollscheid et al. 2016). pcDNA CMV-10 3xFlag-OFD1 full length (1-1012), a (1-276), b (277-663), c (664-1012), and ΔLIR (mutation of the LIR domain EKYMKI to EKAMKA) were kindly provided by Brunella Franco (TIGEM, Napoli). pEGFP-OFD1 full length (FL) construct was generated by subcloning the OFD1 gene from the pcDNA CMV-10 3xFlag-OFD1 construct into the pEGFP_C1 vector with SmaI/BamHI restriction enzymes. pLVX-GFP-OFD1 lentiviral construct was generated using the Infusion HD cloning system (Takara Clontech) following manufacturer’s instructions. In brief, EGFP-OFD1 sequence was amplified by PCR using primers that anneal on the GFP template and are complementary to pLVX-Puro vector (forward: 5’- CTCAAGCTTCGAATTCATGGTGAGCAAGGGCGAG-3’; reverse: 5’- TAGAGTCGCGGGATCCATCAGTTATCTAGATCCGGTGG-3’), followed by EcoRI/BamHI vector linearization and In-fusion reaction. pSLIK-NEO myosin VI shRNA was generated with Gateway™ LR Clonase™ II Enzyme mix (ThermoFisher Scientific) by subcloning a nucleotide sequence targeting myosin VI (5’- AGTAATTCAGCACAATATTCCAA - 3’) into a pENTR vector, followed by recombination into pSLIK-NEO empty vector.

All the other truncated constructs were engineered by site-directed mutagenesis or recombinant PCR and sequence-verified. Details are available upon request.

### Antibodies

| Primary antibodies |  |  |  |  |
| --- | --- | --- | --- | --- |
| Epitope | Species | Supplier | Identifier | Application |
| 53BP1 | Rabbit | Abcam | ab36823 | WB 1:1000 |
| Acetylated tubulin | Mouse | Sigma | #T7451 | IF 1:1000 |
| Acetylated tubulin | Mouse | Sigma | 6-11B-1 | IF 1:1000 |
| Centrin1 | Mouse | Millipore | 04-1624 | IF 1:1000 |
| Cep135 | Rabbit | Abcam | ab75005 | IF 1:1000 |
| Cep164 | Goat | Santa-Cruz Biotechnology | sc-240226 | IF 1:700 |
| FLAG | Mouse | Sigma | F3165 | WB 1:5000 |
| GAPDH | Mouse | Santa-Cruz Biotechnology | Sc-32233 | WB 1:5000 |
| GFP | Rabbit | Sigma | G1544 | WB 1:5000 |
| **GFP** | Mouse | ThermoFisher Scientific | A-11120 | PLA 1:2000 |
| **GST** | Rabbit | generated in house |  | IP |
| **LC3** | Rabbit | Cell Signaling | 12741 | WB 1:1000 |
| **Myosin VI** | Mouse | Sigma | MUD-19 | WB 1:1000 |
| **Myosin VI** | Rabbit | generated by EUROGENTEC S.A., purified by Cogentech | 1295 | IP |
| **Myosin VI** | Rabbit | generated by EUROGENTEC S.A., purified by Cogentech | 1296 | WB 1:2000; IF 1:400 |
| **OFD1** | Rabbit | Sigma | HPA031103 | IF 1:500; PLA 1:2000 |
| **OFD1** | Rabbit | Kindly provided by Brunella Franco (TIGEM, Napoli) (Giorgio et al. 2007) |  | WB 1:100 |
| **p21** | Rabbit | Cell Signaling | 2947 | WB 1:1000 |
| **p53 (DO-1)** | Mouse | Cell Signaling | 18032 | WB 1:1000 |
| **p62** | Rabbit | Enzo Lifesciences | BML_PW9860 | WB 1:1000 |
| **PCM1** | Mouse | Sigma | SAB1406228 | IF 1:200 |
| **Pericentrin** | Mouse | Abcam | 28144 | IF 1:200 |

| **Secondary antibodies** |  |  |  |  |
| --- | --- | --- | --- | --- |
| **Antibody** | **Species** | **Supplier** | **Catalog number** | **Application** |
| **Anti-mouse IgG HRP** | Goat | Bio-Rad | 1721011 | WB 1:5000 |
| **Anti-rabbit IgG HRP** | Goat | Bio-Rad | 1706515 | WB 1:5000 |
| **Anti-mouse Alexa488** | Donkey | ThermoFisher Scientific | A21202 | IF 1:400 |
| **Anti-mouse Cy3** | Donkey | Jackson Lab | 715-165-150 | IF 1:400 |
| **Anti-mouse Alexa488** | Goat | Molecular Probes | A11001 | IF 1:500 |
| **Anti-rabbit Alexa555** | Donkey | Molecular Probes | A31572 | IF 1:500 |
| **Anti-rabbit Alexa488** | Donkey | ThermoFisher Scientific | A21206 | IF 1:400 |
| **Anti-rabbit Cy3** | Donkey | Jackson Lab | 711-165-152 | IF 1:400 |
| **Anti-rabbit Alexa647** | Donkey | ThermoFisher Scientific | A31573 | IF 1:400 |
| **Anti-goat Alexa647** | Donkey | ThermoFisher Scientific | A21447 | IF 1:400 |

Commercial antibodies were validated by the manufacturer. The home-made anti-myosin VI was previously described (Wollscheid et al. 2016).

### Cell lines

hTERT-RPE1 cells (ATCC) were maintained in Dulbecco’s Modified Eagle Medium: Nutrient Mixture F-12 (DMEM/F12, Gibco), supplemented with 10% fetal bovine serum (FBS), 2 mM L-glutamine, 0.5 mM Na-Pyruvate, 15mM Hepes pH 7.5. HEK293T (ICLC) and A549 (NCI-60) cells were maintained in DMEM, supplemented with 10% FBS and 2 mM L-glutamine.

hTERT-RPE1 p53 knock-out (KO) and 53BP1 KO were kindly provided by Luca Fava (University of Trento, Trento). In brief, hTERT-RPE1 cells were transduced with Lenti-CRISPR-V2 targeting coding exons of the genes of interest, selected with puromycin, and single clones were characterized through sequencing of the targeted genomic region (Burigotto et al. 2021).

hTERT-RPE1 with stable expression of centrin1-dTomato were kindly provided by Francesca Farina (IRTSV, Grenoble). In brief, hTERT-RPE1 cells were transfected with pdTomato-centrin1 construct (Farina et al. 2016) and selected with geneticin. dTomato-positive cells were selected by fluorescence-activated cell sorting (FACS). To generate the hTERT-RPE1 centrin1-dTomato GFP-OFD1 cell line used for fluorescence recovery after photobleaching (FRAP) experiments, hTERT-RPE1 centrin1-dTomato were transduced with the pLVX-GFP-OFD1 lentiviral construct. Two weeks after transduction, GFP- and dTomato-positive cells were selected by FACS.

To generate A549 myosin VI KO cell line through CRISPR/Cas9, A549 cells were transiently transfected with pD1301-AD vector (prepared by ATUM, atum.bio), containing Cas9, a GFP reporter and a guide RNA (gRNA) targeting exon 2 of myosin VI (gRNA sequence: 5’-GTTCAATTGTTAAGCTGTCG-3’, designed using ATUM’s design tool). GFP-positive cells were FACS-sorted, single cell clones were screened for myosin VI deletion through immunoblot (IB) and IP, and the genomic mutations were characterized by PCR amplification and sequencing of the targeted genomic region. Clone 13S34 that was selected for our experiments have two different mutant alleles: one has an insertion of 2 nucleotides while the other has a deletion of 19 nucleotides. In both cases the frameshift caused stop codons, generating truncated proteins spanning the first 30 or 38 amino acids, respectively.

All cell lines were authenticated by STR profiling (StemElite ID System, Promega) and tested for mycoplasma using PCR and biochemical test (MycoAlert, Lonza).

### siRNA transfection, shRNA expression

Transient knock-downs (KD) were performed using Stealth siRNA oligonucleotides from Thermo Fischer Scientific (Waltham, MA, USA). Cells were transfected twice using RNAiMax (Invitrogen), first in suspension and the following day in adhesion. The following siRNAs were used: myosin VI #1 (when not specified, this siRNA is used to deplete myosin VI):

5’ - GAGGCUGCACUAGAUACUUUGCUAA-3’

myosin VI #2: 5’-GAGCCTTTGCCATGGTACTTAGGTA-3’

OFD1: 5’-GAGAAUGAAGUGUACUGCAAUCCAA-3’

p53: 5’-CCAGUGGUAAUCUACUGGGACGGAA-3’.

hTERT-RPE1 cells with doxycycline-inducible shRNA for myosin VI were generated by transducing the cells with pSLIK-NEO myosin VI shRNA and selection with neomycine. To obtain myosin VI depletion, the expression of the shRNA was induced with 0.5 µg/ml doxycycline for 10 days.

### Cell lysis, SDS-PAGE, and immunoblotting (IB)

Cells were pelleted and the dry pellet was either frozen at −80°C or directly processed. Cell pellets were lysed in JS buffer (50 mM Hepes, pH 7.5, 150 mM NaCl, 1.5mM MgCl_2_, 5mM EGTA, 10% glycerol, and 1% Triton X-100) supplemented with 20 mM sodium pyrophosphate, pH 7.5, 250 mM sodium fluoride, 2 mM PMSF, 10 mM sodium orthovanadate, and protease inhibitors (Calbiochem) and incubated for 20 minutes on ice. Lysates were cleared by centrifugation at 15,000 rpm for 20 minutes at 4°C. Protein concentration was measured by the Bradford assay (Biorad) following the manufacturer’s instructions. Proteins were denatured by adding Laemmli Buffer to 1X and by boiling at 95°C for 5 minutes. Proteins were then separated on precast gradient gels (4–20% TGX precast gel, Bio-Rad) by SDS-PAGE and transferred to nitrocellulose membranes by Transblot Turbo (BIO-RAD). Membranes were blocked for 1 hour (or overnight) in 5% milk in TBS supplemented with 0.1% Tween (TBS-T). The primary antibodies were diluted in TBS-T 5% milk, and incubated onto the membranes for 1 hour at room temperature (RT), or overnight at 4°C. After washes, the membranes were incubated with the appropriate horseradish peroxidase (HRP)-conjugated anti-mouse or anti-rabbit secondary antibodies (GE Healthcare), subsequently detected with ECL (GE Healthcare). Immunoblots were visualized using films (GE Healthcare).

### Protein expression and purification

GST fusion proteins were expressed in E. coli Bl21 (DE3) Rosetta at 18°C for 16 hours after induction with 0,5 mM IPTG at an OD_600_ of 0.6. Cell pellets were resuspended in lysis buffer (50 mM Na-HEPES, pH 7.5, 250 mM NaCl, 1 mM EDTA, 5% Glycerol, 0.1% NP40, Protease Inhibitor Cocktail set III, Calbiochem, and 0.1 mM PMSF). Sonicated lysates were cleared by centrifugation at 16,000 rpm for 30 minutes. Supernatants were incubated with 1 ml of glutathione-sepharose-beads (GE Healthcare) per litre of bacterial culture. After 2 hours at 4°C, the beads were washed with lysis buffer, high salt buffer (50 mM Tris, pH 8, 1M NaCl, 1 mM EDTA, 1 mM DTT, and 5% glycerol) and equilibrated in storage buffer (20 mM Tris, pH 8, 150 mM NaCl, 1 mM EDTA, 1 mM DTT, and 5% glycerol).

### Biochemical experiments

For co-immunoprecipitation (co-IP) HEK-293T cells were transfected with the indicated constructs. After 48 hours, the cells were lysed in JS buffer and incubated for 20 minutes on ice. Lysates were cleared by centrifugation at 15,000 rpm for 20 minutes at 4°C. Anti-Flag IP was performed by incubating 1 mg lysate with anti-Flag M2 conjugated beads (Sigma) for 2 hours at 4°C. For anti-myosin VI IP, 1 mg lysate was incubated with anti-myosin VI antibodies (1295 or 1296) or anti-GST rabbit antibody as negative control. After 2 hours of incubation at 4°C, protein A sepharose beads were added to the IP and the mixture was incubated for an additional hour. Precipitated immunocomplexes were washed, loaded on a precast gradient gel (4–20% TGX precast gel, Bio-Rad) and analysed by IB using specific antibodies.

For pull-down experiments, 1 μM of GST-fusion proteins immobilised onto GSH beads were incubated for 2 hours at 4°C in JS buffer with 500 µg of transfected HEK293T cellular lysate. Detection was performed by IB. Ponceau-stained membrane was used to show loading of GST-fusion proteins.

### Liquid chromatography–tandem MS (LC–MS/MS) analysis

To identify myosin VI interactors, anti-myosin VI IP was performed in HeLa, MDA-MB-231, MCF10A, and Caco-2 cells grown in confluent conditions. 3 mg of lysates were incubated with anti-myosin VI antibody (1295) or a rabbit control antibody as negative control. After 2 hours of incubation at 4°C, protein A sepharose beads were added to the IP and the mixture was incubated for an additional hour. Precipitated immunocomplexes were washed, loaded on a precast gradient gel (4–20% TGX precast gel, Bio-Rad) and stained with colloidal blue (Colloidal Blue Staining Kit, Invitrogen).

Slices of interest were cut from gels and trypsinized as previously described by Shevchenko and colleagues (Shevchenko et al. 1996). Peptides were desalted as described by Rappsilber and colleagues (Rappsilber, Ishihama, and Mann 2003), dried in a Speed-Vac and resuspended in 10 µL of solvent A (2% ACN, 0.1% formic acid). 3 μL were injected on a quadrupole Orbitrap Q-Exactive mass spectrometer (Thermo Scientific) coupled with an UHPLC Easy-nLC 1000 (Thermo Scientific), with a 25 cm fused-silica emitter of 75 μm inner diameter. Columns were packed in-house with ReproSil-Pur C18-AQ beads (Dr. Maisch Gmbh, Ammerbuch, Germany), 1.9 μm of diameter, using a high-pressure bomb loader (Proxeon, Odense, Denmark). Peptide separation was achieved with a linear gradient from 95% solvent A (2% ACN, 0.1% formic acid) to 40% solvent B (80% acetonitrile, 0.1% formic acid) over 30 minutes and from 40% to 100% solvent B for 2 minutes at a constant flow rate of 0.25 μL/minute, with a single run time of 33 minutes. MS data were acquired using a data-dependent top 12 method, and the survey full scan MS spectra (300–1750 Th) were acquired in the Orbitrap with 70000 resolution, AGC target 1e6, IT 120 ms. For HCD spectra, the resolution was set to 35000, AGC target 1e5, IT 120 ms; 25% normalised collision energy and isolation width of 3.0 m/z.

For protein identification, the raw data were processed using Proteome Discoverer (version 1.4.0.288, Thermo Fischer Scientific). MS2 spectra were searched with Mascot engine against uniprot_human_20150401 database (90411 entries), with the following parameters: enzyme Trypsin, maximum missed cleavage 2, fixed modification carbamidomethylation (C), variable modification oxidation (M) and protein N-terminal acetylation, peptide tolerance 10 ppm, MS/MS tolerance 20 mmu. Peptide Spectral Matches (PSM) were filtered using percolator based on q-values at a 0.01 FDR (high confidence). Proteins were considered identified with 2 unique high confident peptides (Kall et al. 2007). Scaffold (version Scaffold_4.3.4, Proteome Software Inc., Portland, OR) was used to validate MS/MS based peptide and protein identifications. Peptide identifications were accepted if they could be established at a probability greater than 95.0% by the Peptide Prophet algorithm (Keller et al. 2002) with Scaffold delta-mass correction. Protein identifications were accepted if they could be established at a probability greater than 99.0% and contained at least 2 identified peptides. Protein probabilities were assigned by the Protein Prophet algorithm (Nesvizhskii et al. 2003). Proteins that contained similar peptides and that could not be differentiated based on MS/MS analysis alone were grouped to satisfy the principles of parsimony. Proteins sharing significant peptide evidence were grouped into clusters.

### Immunofluorescence (IF)

Cells were grown on coverslips and fixed with cold 100% methanol at −20°C for 10 minutes. The coverslips were incubated in PBS with 10% FBS for 30 minutes for blocking, followed by incubation with primary antibodies (overnight at 4°C) and then secondary antibodies (1 hour at RT) in PBS with 10% FBS. Incubation with DAPI (Sigma-Aldrich, cat. D9542) for 10 minutes was performed to stain the nuclei. The coverslips were mounted on glass slides using Mowiol Mounting Medium (Calbiochem) or Dako Faramount Aqueous Mounting Medium (s3025, DAKO). Images were acquired using a GE HealthCare Deltavision OMX system, equipped with 2 PCO Edge 5.5 sCMOS cameras, using a 60x 1.42 NA Oil immersion objective, or using a Deltavision Elite system (GE Healthcare) equipped with a IX71 microscope (Olympus), a sCMOS camera, using a 60x PlanApo 1.42 NA oil immersion objective and driven by softWoRx version 7.0.0. Images were acquired as a z-series (0.2-µm z interval in a range of 4 µm) and deconvolution was performed using softWorx software. The images are presented as maximum-intensity projections and were prepared using ImageJ (National Institutes of Health).

### OFD1, cep164 and PCM1 fluorescence intensity analysis

For OFD1 and cep164 fluorescence intensity quantifications at the centrioles, hTERT-RPE1 cells were treated with 6 µg/ml nocodazole (M1404, Sigma) for 1 hour to depolymerize the microtubules and thus remove the centriolar satellites. The removal of the satellites is required for the quantification of the centriolar pool of OFD1, which is otherwise indistinguishable from the satellite pool. Z-stack images (0.2 µm intervals) of 20 sections were acquired as detailed above. Centrioles were considered for quantification when paired signals of centrin and OFD1 were observed. The presence cep164 staining was used to identify mother centrioles. Centrioles were excluded from the analysis if residual OFD1 satellite staining was visible in the vicinity of the centrioles. The integrated intensity of a circular 1×1 µm area around the specific OFD1 and cep164 signals were recorded in the sum projected images using ImageJ software (National Institutes of Health). Integrated density was corrected for background intensity signal with the following formula: corrected total fluorescence = integrated density – (area of selected region*mean intensity of background). The number of cells analysed and of experiments performed are detailed in the figure legends.

For centriolar satellite intensity quantifications, staining of PCM1 was performed in combination with a centriole marker (cep135 or cep164). The centriole marker was used to determine the centre of a circular 3×3 µm area, where the PCM1 signals were recorded in the sum projected images. For quantification of OFD1 intensity at the centriolar satellites, OFD1 staining was performed in combination with PCM1 and cep164. In order to exclude the contribution of the centriolar pool of OFD1 from the analysis, OFD1 intensity was recorded only in the area covered by PCM1 in the sum projected images. Corrected total fluorescence was calculated as detailed above.

To induce p53 activation, hTERT-RPE1 cells were incubated with 10 µM Nutlin-3 (N6287, Sigma) for 24 hours before fixation and IF.

### Statistical analysis

Statistical analyses were performed with Prism (GraphPad software). Unless differently specified, all the statistical significance calculations were determined using either unpaired Student’s T or ANOVA test, or the non-parametric Mann Whitney or Kruskal-Wallis test, after assessing the normal distribution of the sample with Normal (Gaussian) distribution test. Sample sizes are indicated in the figure legends and were chosen arbitrarily with no inclusion and exclusion criteria. The investigators were not blind to the group allocation during the experiments and data analyses.

### Proximity ligation assay (PLA)

For GFP-myosin VI overexpression, hTERT-RPE1 cells were transfected with pEGFP-C1 myosinVI_short_ FL using Lipofectamine 2000 reagent (Invitrogen), following manufacturer’s instruction. Fixation was performed 48 hours after transfection with 100% MeOH at −20°C for 10 minutes. The PLA was performed with the Duolink™ In Situ Orange Starter Kit (Sigma, DUO92102) according to manufacturer’s instructions. Primary antibodies used for PLA include mouse anti-GFP (1:2000; ThermoFisher Scientific, A11120) and rabbit anti-OFD1 (1:2000; Sigma, HPA031103). Secondary anti-mouse MINUS and anti-rabbit PLUS probes were used. As negative controls for the PLA signal, the secondary antibodies were used without previous primary antibody incubation or with the single primary antibody. After the PLA procedure, counterstaining with anti-mouse A488 and anti-rabbit A647 was performed to identify GFP-positive cells and to localise OFD1. Confocal microscopy was performed on a Leica TCS SP5 laser confocal scanner mounted on a Leica DMI 6000B inverted microscope equipped with motorised stage. The images were acquired with an HCX PL APO 63X/1.4NA oil immersion objective using 405 nm, 488 nm, 568 nm, and 647 nm laser lines. Leica LAS AF software was used for all acquisitions.

### Fluorescence recovery after photobleaching (FRAP)

hTERT-RPE1 centrin1-dTomato GFP-OFD1 cells were transfected with myosin VI siRNA and plated on MatTek glass bottom dishes (P35G-1.5-14-C, MatTek Life Sciences). FRAP experiments were performed on the UltraVIEW VoX spinning-disk confocal system (PerkinElmer) equipped with an EclipseTi inverted microscope (Nikon), provided with an integrated FRAP PhotoKinesis unit (PerkinElmer), a Hamamatsu CCD camera (C9100-50), a Okolab cage incubator, and driven by Volocity software (Improvision; Perkin Elmer). Photobleaching was achieved on a square region of 3×3 µm by using the 488 nm laser at the maximum output to bleach the GFP signal. Initially, 10 images with a 400 ms time-frame were acquired to determine the levels of pre-bleach fluorescence. Images were acquired through a 60X oil-immersion objective (Nikon Plan Apo VC, NA 1.4) with the following time-frame: every second for the first 60 seconds, every 2 seconds for the following 60 seconds and every 5 seconds for the following 120 seconds.

A custom ImageJ macro and a set of functions written in Matlab software were used to analyse the recovery curves. ImageJ was used to measure the mean intensity value over time in the bleached area and the background over time in an area in the field without cells. StackReg ImageJ Plugin was used to align the bleached area over time. The photobleaching recovery curves were then normalised to the pre-bleaching mean intensity values after background correction using Matlab. Matlab was then used to fit the first 150 seconds after photobleaching of the recovery curves with a one phase exponential recovery function with f0 fixed as the first value after photobleaching:

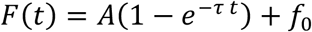

Half maximum time (t_1/2_) was then evaluated as:

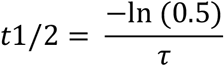

and Mobile Fraction as:

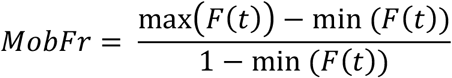

### Structured illumination microscopy (SIM)

hTERT-RPE1 control and myosin VI-depleted cells were plated on 13 mm coverslips, thickness 1.5 mm, Boroslicate glass (631-0150, VWR). Four days after transfection, cells were incubated for 1 hour on ice in PBS to induce the depolymerisation of cytoplasmic microtubules, and were subsequently stained with anti-acetylated tubulin and anti-OFD1 antibodies (for more details, IF staining section). SIM Images were acquired using a GE HealthCare Deltavision OMX system, equipped with 2 PCO Edge 5.5 sCMOS cameras, using a 60x 1.42NA Oil immersion objective. Images were deconvolved and reconstructed with Applied Precision’s softWorx software. The images are presented as maximum-intensity projections and were prepared using ImageJ (National Institutes of Health).

### Primary cilium assay

hTERT-RPE1 p53 KO cells were transfected with myosin VI or OFD1 siRNAs and plated on coverslips coated with 0.2% gelatine. Four days after transfection, cells were serum starved in medium with 0% serum (DMEM/F12 supplemented with 2 mM L-glutamine, 0.5 mM Na-Pyruvate and 15 mM Hepes, pH 7.5) for 24 hours to allow cilium assembly. Microtubule depolymerisation was induced by incubating the cells in PBS at 4°C for 1 hour before Me-OH fixation. Subsequently, cell lines were stained with DAPI, anti-acetylated tubulin and anti-cep135 primary antibodies. Z-stack images (0.2 µm intervals) of 20 sections were acquired using a Deltavision Elite system (GE Healthcare) equipped with a IX71 microscope (Olympus), a sCMOS camera, using a 60x PlanApo 1.42 NA oil immersion objective and driven by softWoRx version 7.0.0. The number of cells displaying a primary cilium was determined using anti-acetylated tubulin staining, and centrioles were identified through the double cep135/acetylated tubulin staining.

### Correlative Light Electron Microscopy (CLEM)

hTERT-RPE1 cells were transfected with the indicated siRNA 96 hours before fixation and plated on gridded MatTek glass bottom dishes (P35G-1.5-14-CGRD, MatTek Life Sciences). Cells were fixed and stained as described by Mironov & Beznoussenko (Mironov and Beznoussenko 2013), using anti-pericentrin antibody to identify the position of the centrosomes. Images were acquired on a XZ plane to calculate the distance between the glass bottom and the centrosome, using a Leica TCS SP5 laser confocal scanner mounted on a Leica DMI 6000B inverted microscope equipped with a HCX PL APO 63X/1.4NA oil immersion objective using 488 nm laser line. Leica LAS AF software was used for all acquisitions. Samples were then fixed with 2.5% GA in 0.2 M Cacodylate buffer, pH 7.2, and embedded in resin (Beznoussenko and Mironov 2015). The samples embedded in resins were sectioned with diamond knife (Diatome, Switzerland) using Leica EM UC7 ultramicrotome. Sections (50-60 nm) were analysed with a Tecnai 20 High Voltage EM (FEI, The Netherlands) operating at 200 kV.

For immunogold staining of myosin VI, A549 wild-type and myosin VI KO cells were plated on gridded MatTek glass bottom dishes and treated as detailed/described above for CLEM. Then, cells were stained with anti-myosin VI 1296 antibody, incubated with Nanogold®-Fab’ Goat anti-Rabbit IgG (#2004, Nanoprobes). GoldEnhance™ (EM #2113, Nanoprobes) was used to enhance the signal from nanogold particles.

### Growth curve

Cells were transfected with the indicated siRNAs. After 24 hours, 10,000 cells were plated on a 6-well plate (day 0). From day 3 to 7 after plating, the cells were detached and counted using Beckman Multisizer 3 Coulter Counter. Cell counts were plotted to display the growth curve. During the experiments, the cells were split upon reaching 70% confluence to avoid slowdown of proliferation due to contact inhibition. Cell dilution was taken into consideration for the total cell count.

### Flow cytometry analysis of cell cycle profile

Myosin VI shRNA expression was induced with 0.5 µg/ml doxycycline for 10 days. The cells were then trypsinized and washed once in PBS. After centrifugation, cell pellets were resuspended in 250 µl of 4°C-cold PBS and fixed by adding 750 µl of 100% Et-OH (−20°C) dropwise while vortexing. The cells were left in the fixative for 1 hour on ice, washed in PBS-1% BSA, and resuspended in 1 ml PI (50 µg/ml) and RNAse (250 µg/ml). After incubation at 4°C overnight, flow cytometry was performed for cell cycle analysis. Sample acquisition was performed with FACSCanto II (Beckton Dickinson). Analysis of cell cycle distribution was performed with ModFitLT V3.1 software.

### Senescence-associated-β-galactosidase (SA-β-gal) assay

Cells were grown on coverslips, washed in PBS and fixed with 4% PFA for 10 minutes. Then, the cells were incubated with SA-β-gal staining solution containing 1 mg 5-bromo-4-chloro-3-indolyl beta-D galactopyranoside (X-Gal) per ml, 40 mM citric acid/sodium phosphate pH 6.0, 5 mM potassium ferrocyanide, 5 mM potassium ferricyanide, 150 mM NaCl, and 2 mM MgCl2. After incubation at 37°C overnight, the cells were washed with PBS and mounted on glass slides using Mowiol Mounting Medium. Images were acquired using a digital camera connected to a white-light microscope. SA-β-gal activity was detected in senescent cells as local perinuclear blue precipitate.

